# Multi-environment Genome Wide Association Studies of Yield Traits in Common Bean (*Phaseolus vulgaris* L.) × Tepary Bean (*P. acutifolius* A. Gray) Interspecific Advanced Lines at the Humid and Dry Colombian Caribbean Subregions

**DOI:** 10.1101/2022.08.03.502649

**Authors:** Felipe López-Hernández, Esteban Burbano-Erazo, Rommel Igor León-Pacheco, Carina Cecilia Cordero-Cordero, Diego F. Villanueva-Mejía, Adriana Patricia Tofiño-Rivera, Andrés J. Cortés

## Abstract

Genome Wide Associations Studies GWAS are a powerful strategy for the exploration adaptive genetic variation to drought stress in advanced lines in common bean with interspecific genotypes, yet they still lack behind in the use of arid multi-environments as the subregions of the Colombian Caribbean. In order to bridge this gap, we couple an advanced genotypes panel integrated with Common Bean (*Phaseolus vulgaris* L.) × Tepary Bean (*P. acutifolius* A. Gray) interspecific lines with GWAS algorithms to identify novel sources of drought tolerance across the subregions of Colombian Caribbean. One of the most important challenges in agriculture is to achieve food security in environments vulnerable to climate change which worsens with the passing of the years. The common bean, a key product of the food basket of vulnerable regions of the Caribbean is affected by the reduction in yield under drought stress. A total of 87 advanced accessions with interspecific lines were genotyped by sequencing (GBS), leading to the discovery of 15,645 single-nucleotide polymorphism (SNP) markers. Five yield traits were developed for each accession and inputted in GWAS algorithms (*i.e.* FarmCPU, and BLINK) to identify putative associated loci in drought stress. Best-fit models revealed 47 significantly associated alleles distributed in all 11 common bean chromosomes. Flanking candidate genes were identified using 1-kb genomic windows centered in each associated SNP marker. A pathways enriched analysis was carried out using the mapped output in the GWAS step for each yield traits indices. Some of these genes were directly linked to response mechanisms of drought stress to level morphological, physiological, metabolic, signal transduction, and fatty acid and phospholipid metabolism. This work offers putative associated loci for marker-assisted and genomic selection for drought tolerance in common bean. It also demonstrates that it is feasible to identify genome-wide associations with an interspecific panel of genotypes and modern GWAS algorithms in multiples environments.

## INTRODUCTION

One of the most important challenges in agriculture is to achieve food security in environments vulnerable to climate change. It is estimated that by 2020, Latin America and the Caribbean presented levels of undernourishment of 47.7 million people, and a projection of 67 million by 2030, the above without considering the repercussions of the COVID-19 pandemic (FAO, 2020). Legumes as beans are one of the main nutritional contributions for the communities of Latin America, Africa and Asia, being nutritionally rich due to their high content of protein, folic acid, iron, dietary fiber and complex carbohydrates (Sgarbieri and Whitaker, 1982). The common bean (*Phaseolus vulgaris* L.) is one of the most produced legumes with ~27 million tons worldwide, with China and America being the main producers (FAO, 2018). Faced with a climate change scenario, the global average temperature will increase by values close to the 1.5°C threshold established in the *Paris agreement* (Shukla et al., 2019) with projections of a decrease in average precipitation for the Caribbean Colombian (Teichmann et al., 2013; Sánchez et al., 2015). The low levels of precipitation and the increase in temperature in the Caribbean will limit the productivity of beans and other species in the regional food basket, that in addition to being an important component of food security in the region, are part of the cultural heritage of some indigenous communities (Tofiño Rivera et al.). However, breeding efforts have important challenges to obtain genotypes with good adaptability to adverse abiotic conditions (Beebe et al., 2013) due to the polygenic nature of the tolerant phenotypes.

A strategy to develop cultivars with tolerance to heat and drought stress requires the efficiently using genetic resources from other closely genepools related species adapted to these abiotic conditions (Buitrago-Bitar et al., 2021). The Tepary bean (*P. acutifolius* A. Gray) is an annual autogamous bean native of the northwest Mexico (Jiri et al., 2017; Mhlaba et al., 2018), domesticated near the arid border with the USA, and adapted to hot (Moghaddam et al., 2021) and dry environments (Muñoz et al., 2006; Mwale et al., 2020). Thus, the Tepary bean is the most heat tolerant of the genus *Phaseolus* due to this natural adaptation. Nowadays, the Tepary bean is limited as modern crop compared with the more susceptible but commercially accepted, the common bean. In this sense, authors as (Burbano-Erazo et al., 2021) have suggest the using the Tepary could serve as donor of alleles to boost drought and heat tolerance in common bean. Previous efforts have set the basis to explore the potential of the Tepary bean as an exotic donor, for instance, the common bean has been backcrossed with Tepary donors with a relatively good rate of viability (Belivanis and Doré, 1986; Souter et al., 2017). Yet, these efforts were unable to pyramid target alleles for drought tolerance, possibly due to its complex quantitative genetic inheritance (Muñoz et al., 2006)

To improve our understanding of the genetic basis of quantitative traits, identification of as many causal loci as possible is imperative (Cortés and Blair, 2018). The exploration of the genetic basis of quantitative traits in crops has been accelerated by modern genomic strategies such as Genome Wide Associations Studies (GWAS) and Genome–Environment Associations Studies (GEAS) (Frank et al., 2016). For instance, the exploring of genetic architecture of abiotic stress in common beans as drought (Cortés and Blair, 2018) and heat (López-Hernández and Cortés, 2019) under GEA paradigm. Also, several GWAS works have studied breeding attributes in beans such as: agronomic traits (Diaz et al., 2020), mineral concentration (Wu et al., 2020), or aluminum toxicity tolerance (Ambachew and Blair, 2021).

Several plant breeders have explored the use of interspecific panel of genotypes using GWAS to improve the reconstruction of the genetic basis of quantitative traits. In the tree context, some authors have used hybrids genotypes in GWAS for phytochemical, morphological and growth traits in hybrid *Populus* (Bresadola et al., 2019), and biotic resistance in hybrid grape (Vasylyk et al., 2022). In oil palms, authors as (Osorio-Guarín et al., 2019) have implemented hybrid *Elaeis* in GWAS for morphological and yield-related traits. Similarly, in cotton, plant scientistic have integrated hybrid genotypes of cotton in GWAS analysis for agronomic traits. In crops, authors have utilized interspecific panel genotypes as: *Amaranthus* for flowering time (Lin et al., 2022); Blueberry (*Vaccinium*) for phenology-related traits and their adaptive variations (Nagasaka et al., 2022); and wheat for grain yield and yield-related traits in drought-stressed (Bhatta et al., 2018). In legumes as beans, relevant to food security, no significant progress has yet been reported on the implementation of interspecific panels in Common bean × Tepary bean using modern GWAS algorithms (Wang and Zhang, 2021) that promote improvement plans by exploring potential alleles associated with adaptation to hot and dry regions such as the Colombian Caribbean, currently with food security problems.

Nowadays, there is a lack of knowledge on how a multi-environment Genome Wide Association Studies work under the advanced genotypes panel integrated with common bean (*Phaseolus vulgaris* L.) × tepary bean (*P. acutifolius* A. Gray) interspecific lines in the humid and dry Colombian Caribbean subregions with the yield traits. Additionally, there is an urgent need to identify loci linked to drought and heat stress tolerance in common bean germplasm collections with interspecific lines, which would aid the development of common bean varieties resistant to high drought or temperature. Therefore, for this study, we set the following objectives: (1) synthetize yield indices in order to estimate drought and heat stress tolerance in wild common bean germplasm collections with interspecific lines, which would allow identifying tolerant accessions; (2) explore the utility of the common bean germplasm collections with interspecific lines under the modern GWAS models (FarmCPU, and BLINK); and (3) implement GWAS models in the Caribbean subregions in order to capture adaptive genetic variation to drought stress, candidate to be integrated into common bean breeding programs for this localities. This first exploration of the local adaptation of common bean germplasm collections with interspecific lines in the Caribbean environments which to will ultimately offer putative associated loci for marker-assisted and genomic selection strategies by using a combination of various well-chosen agronomic indices and GWAS algorithms, while testing the utility of the latter under a interespecific paradigm.

## MATERIALS AND METHODS

### Plant material

The panel of 87 genotypes used in this GWAS study is composed of 67 interspecific lines between the common bean (*P. vulgaris*) and Tepary bean (*P. acutifolius),* and 19 advanced genotypes bred to high temperature and drought conditions by the bean program of the Alliance Bioversity–CIAT (International Center for Tropical Agriculture) and transferred to AGROSAVIA after ATM subscription. In addition, the genotype G40001 (*P. acutifolius*) was used as control. The interspecific lines correspond to the third generation (and beyond, detailed pedigree in Table S1) evaluated for the first time at four localities in the humid and dry Colombian Caribbean subregions (Burbano-Erazo et al., 2021).

### Multi-Locality Field Trials

During the crop cycle of July–October 2020, the GWAS panel was evaluated at four localities in the humid and dry Colombian Caribbean subregions corresponding to the following AGROSAVIA’s research stations: Motilonia Research Station ([10°00’01.2”N, 73°15’22.4”W] Codazzi, municipality in the state of Cesar), Caribia Research Station ([10° 47’ 35,4”N, 74° 10’ 49,9”W] Sevilla, municipality in the state of Magdalena), Carmen de Bolívar Research Station ([9°42’50.8”N, 75°06’26.9”W] Carmen de Bolívar, municipality in the state of Bolívar), and Turipaná Research Station ([8°50’27.47”N, 75°48’27.56”W] Cereté, municipality in the state of Córdoba. The research stations Motilonia and Carmen de Bolivar (Mountains and Piedmont, both at more than 100 m.a.s.l.) belong to the dry Caribbean sub-region, while the research station Caribia and Turipaná (Plain, both at less than 20 m.a.s.l.) were representatives of the humid Caribbean sub-region.

With the aim to validate this environmental sub-region, these environmental variables were measured *in situ* in the months from cultivation to harvest: precipitation, temperature and humid relative, suggesting that the research stations Motilonia and Carmen de Bolivar had less precipitation (153.25 mm ± 34.94 and 81.95 mm ± 40.30 respectively) than the research station Caribia and Turipaná with high precipitation (207.733 mm ± 106.80 and 214.85 mm ± 92.62 respectively). The RS with the highest relative humidity was Motilonia and the RS with the lowest relative humidity was Carmen de Bolivar (Table S2). Also, using the Global daily (1km) land surface precipitation based on cloud cover-informed downscaling (Karger et al., 2021), the time series of daily precipitation 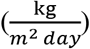 was extracted for the same months from cultivation to harvest from the historical data of year 2003 to year 2016. The data suggested that the two RS with more precipitation were Caribia and Turipaná (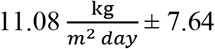 and 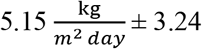 respectively), and the RS with less precipitation were Motilonia and Carmen de Bolivar (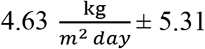 and 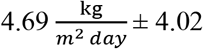 respectively) (Figure S1). The soil characteristics suggested that the RS Carmen de Bolivar had the highest level of pH, P, Ca, and K than the other RS. (Table S2).

### Experimental Design and Phenotyping

The GWAS panel was carried out using a completely randomized block design (CRBD) with three repetitions at each locality. The experimental unit per treatment (genotypes) was a plot of four m^2^ with a spatial arrangement of one row spaced at 0.8 m and 0.25 m between plants (13 plants per genotype). Missing genotypes were indicative of mal-adaptation at each locality. The standard yield traits (Blair et al., 2012; Burbano-Erazo et al., 2021) in common bean measured at the end of the cycle at each locality were, NS: average number of seeds per pod, NP: number of pods per plant, YLP: yield per plant (g/plant), SB: seed weight (g), and VB: vegetative biomass (g).

### Compilation of Indices of Yield Traits and Statistical Analysis

With the aim of weighing the intra-genotype variability in all yield traits, we proposed an index that ponders the variability in each trait to divide the mean of each genotype between its variance. Thus, high values of the index indicate genotypes with high performance and stable in the field.

The GWAS approaches based on Mixed Lineal Models (MLM) as the algorithms of GAPIT3 (Wang and Zhang, 2021) are known to improve the statistical results to normalize the quantitative traits (Goh and Yap, 2009). In this sense, we carried out the normalization of each trait by means of the automatic transformations Tukey’s Ladder of Powers (Tukey, 1977) using the R-package *rcompanion* (Salvatore Mangiafico, 2022). This algorithm uses Shapiro-Wilk tests iteratively to find at which lambda value the data is closest to normality and transforms it. Then, the normality test used to validate the gaussian distribution was Shapiro-wilk, carried out using the R-package *nortest* (Gross and Ligges, 2015). Later, an analysis of variance between localities (Research Stations) was carried out by means of Welch’s one-way ANOVA using the *ggbetweenstats* function in the R-Package *ggstatsplot* (Patil, 2021).

The Games-Howell *post-hoc* test is another non-parametric approach to compare combinations of groups when there is an imbalance in the number of individuals and homogeneity of variance cannot be assumed. Because this study has different sample sizes per location, a Games-Howell test was implemented using Holm’s P-value adjustment method by the *ggbetweenstats* function in the R-Package *ggstatsplot* (Patil, 2021).

### DNA Extraction and Genotyping-by-Sequencing

The genomic data of the GWAS panel was obtained by means of the Genotyping By Sequencing GBS following to (Elshire et al., 2011). The genomic DNA was extracted from 20 mg of the tissue of leaf collected to 40 days after the plant germination immediately stored in silica gel (Sigma-Aldrich, Alemania). The DNA extraction was carried out using the AGROSAVIA’s *in-house* protocol with steps of maceration with liquid nitrogen, organic compounds (phenol and chloroform) and precipitation with alcohols. The quantification of the extracted DNA was evaluated by means of spectrophotometry using the Nanodrop^®^ 2000 equipment (Thermo Fischer Scientific, United States) and by fluorimetric method using the Qubit^®^ dsDNA HS fluorometer (Life Technologies, Sweden). The enzymatic digestion was carried out using the cutting enzyme *Apek1* (Cortés and Blair, 2018; López-Hernández and Cortés, 2019). Subsequently, DNA libraries were prepared using the NEBNext^®^ Ultra™ II DNA Library Prep Kit for Illumina^®^. Later, the DNA libraries were quantified by the fluorimetric method using the Qubit^®^ dsDNA HS fluorometer. Finally, the concentration and fragment sizes of the DNA libraries were evaluated using the TapeStation 4200 kit (Agilent Technologies, United States) and the High Sensitivity D1000 kit.

### SNP calling

DNA sequences were obtained using the Illumina 2500 Hiseq sequencer (Macrogen, South Korea) in a single direction (single-end) and preliminarily analyzed by the FastQC program (Andrews, 2010) using the Illumina 1.9 coding. In order to clean the DNA sequencing data, the algorithm Trimmomatic (Bolger et al., 2014) was run with the main parameters: ILLUMINACLIP:TruSeq3-SE:2:30:10, window SLIDINGWINDOW:4: 20 and MINLEN:20. Subsequently, a quality analysis of the fastq files was performed using the FastQC program (Andrews, 2010) using Illumina 1.9 coding.

With the aim to identify allelic polymorphism in the GWAS panel, an automatized SNP calling script was constructed using the function *HaplotypeCaller* of the protocol GATK4 (McKenna et al., 2010) with the alignment algorithm BWA (Li, 2013). The genome reference used was the second annotated version of *Phaseolus vulgaris* L. arranged in the platform *Phytozome* http://phytozome.jgi.doe.gov/ with extension of 600Mb and dpeth of ~83,2x (Schmutz et al., 2014b). The mapping statistic was obtained by means of the function *flagstat* from Samtools 1.9 software (Li et al., 2009) in the platform of the Galaxy project 2.0.3 (Afgan et al., 2018). The SNP matrix was filtered by the software Tassel 5.2.78 (Bradbury et al., 2007) using a min depth of 3X and percentage of missing data by loci and by sample of 80%. Finally, the SNP calling script are available in the GitHub page (https://github.com/FelipeLopez2019/SNP-calling-of-KOLFACI-project/blob/main/Kolfaci_Colombia_v4.sh).

### Analysis of Kinship and Population Structure

Using the 15,645 SNP markers, the random and fixed effects were estimated in order to reduce the rate of false positives of each GWAS models (BLINK and FarmCPU). Random effects accounted for kinship relationships, while fixed effects accounted for population structure. Kinship was built by means of the VanRaden algorithm available in the GAPIT (Wang and Zhang, 2021) package of R v. 4.1.2 (R Core Team). On the other hand, the population stratification was explored using the molecular principal component analysis (hereinafter referred to as PC) carried out in GAPIT and *optCluster* R-package (Sekula et al., 2017), and the non-negative matrix factorization algorithm (*snmf* function) in the LEA R-package with 10,000 repetitions, an improvement to admixture analysis (Frichot and François, 2015) optimized by Cross-Entropy.

### Identification of Loci Associated with Yield Traits

The GWAS algorithms FarmCPU and BLINK are known to increase the statistical power while better controlling the false-positive rate, which makes them particularly powerful at further controlling the false -negative rate (Liu et al., 2016; Zhang et al., 2018). The multi-environment GWAS models in advanced common Bean × tepary interspecific lines were obtained from the combination of all yield Traits (number of seeds, number of pods, yield, seed biomass and vegetative biomass). A population stratification method as fixed effects (Principal Components), and a kinship method as random effect (VanRaden) were considered for a total of 40 models (10 for each locality), which were constructed by means of the FarmCPU and BLINK algorithms.

Highly significant associations were determined using a Bonferroni correction of *p*-value at an α = 0.05, which led to a significance threshold of -log10 (3,196 E-6) = 5.4954 for all GWAS models. Therefore, we used the Bonferroni threshold in order to evaluate the rate of false positives by visual interpretation of the Q–Q plots. In addition, a relax threshold of -log10 (3,196 E-4) = 3.495, as previously suggested (Pasam et al., 2012; López-Hernández and Cortés, 2019; Oladzad et al., 2019), since it is documented that the Bonferroni threshold is very restrictive or conservative in MLMs (Joo et al., 2016). Finally, the circular manhattan plots and QQplots were generated by the R-package *RIdeogram* (Hao et al., 2020).

### Identification of Candidate Genes

Putative candidate genes were identified by inspecting conservative flanking sections of 1 kb around each associated locus from all GWAS models. Flanking sections were captured using the common bean reference genome v2.1 (Schmutz et al., 2014a) and the *PhytoMine* tool from the Phytozome v.13 platform (https://phytozome-next.jgi.doe.gov/) where the identified genes were further annotated from the GO (http://geneontology.org/), PFAM (https://pfam.xfam.org/), PANTHER (http://www.pantherdb.org/), KEGG (https://www.genome.jp/kegg/), and Uniprot (https://www.uniprot.org/) databases.

### Pathways Enriched Analysis

Pathway enrichment analysis identifies biological pathways that are enriched in a gene list more than would be expected by chance (Reimand et al., 2019). A pathways enriched analysis was carried out using the mapped output in the GWAS step for each yield traits indices. The software used was the *PhytoMine* tool in the Phytozome v.13 platform employing the The MetaCyc database of metabolic pathways and enzymes (Caspi et al., 2018) (https://metacyc.org/). The p-value was calculated using the Hypergeometric distribution using the number of genes of GWAS, the number of genes in the reference population (*Arabidopsis thaliana),* the number of genes annotated with this item in your list, and the number of genes annotated with item in the reference population (*A. thaliana).* Also, a gene ontology enrichment was carried out without obtaining a significant result.

## RESULTS

GWAS panel suggest a segregant behavior across all localities. With the aim of weighing the intra-genotype variability in all yield traits, we proposed an index that ponders the variability in each trait to divide the mean of each genotype between its variance. All traits index were normalized using the automatic transformations Tukey’s Ladder of Powers (Tukey, 1977) obtaining gaussian distribution in each variable. From the GWAS panel and the genome reference of *Phaseolus vulgaris* L., a raw matrix of allelic variants was obtained with 15,645 SNP markers using the GATK protocol after filtering. The genetic structure and kinship relationships suggested five demographic groups and substructure related to families of genotypes. A total of 47 unique loci in all locations with five yield traits indices were detected. 43 associated markers flanked 90 genes across all chromosomes. Finally, pathways enriched for genes associated in the NP, NS, SB, and VB indices: *Palmitate Biosynthesis II, Stearate Biosynthesis II,* and *Superpathway of Fatty Acid Biosynthesis II.*

### GWAS panel suggest a segregant behavior across all localities

The descriptive analysis suggests that most of the genotypes in the GWAS panel presented a segregant behavior (Figure S2-6) in all yield traits. Also, this segregant behavior was systematic in other field trial (2022-I) using a second panel of interspecific genotypes with same parentals.

### All Indices of yield traits were compiled and normalized for GWAS models

All traits index were normalized using the automatic transformations Tukey’s Ladder of Powers (Tukey, 1977) obtaining gaussian distribution in each variable: number of pods per plant Index Figure 1-A, average number of seeds per pod Index Figure 1-B, yield per plant Index (g/plant) Figure 1-C, seed weight Index (g) Figure 1-D, vegetative biomass Index (g) Figure 1-E. Also, through the Shapiro-Wilk test, the normality was corroborated in all localities (Table S3). Thus, all normalized indices were used in the GWAS algorithms to obtain the genetic base of yield traits.

**Figure 1.**
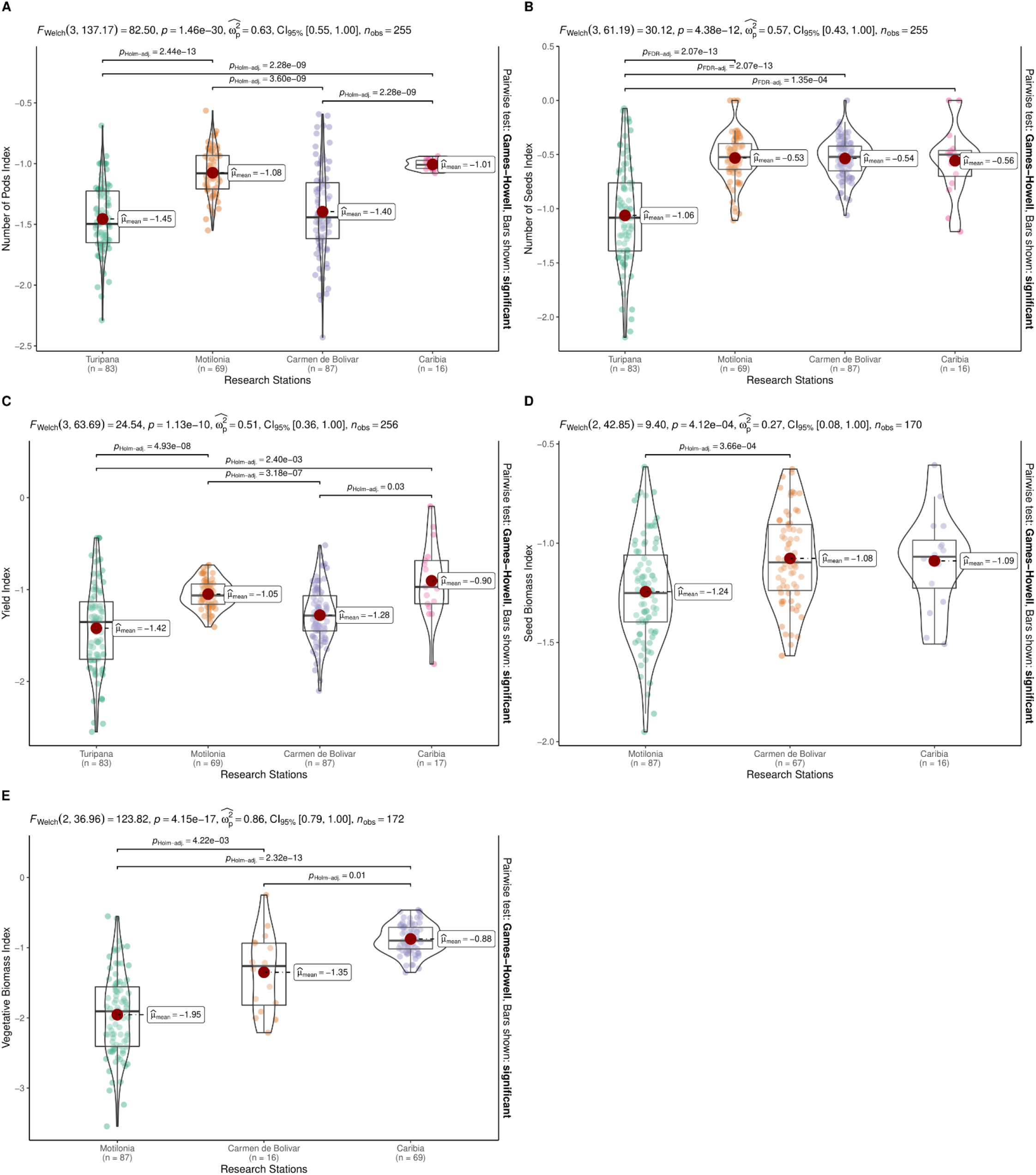
Analysis of variance between locations (Research Stations) by Welch’s one-way ANOVA using the *ggbetweenstats* function in the R-Package *ggstatsplot*. A Games-Howell test was implemented using Holm’s P-value fitting method using the *ggbetweenstats* function in the R-Package *ggstatsplot* (Patil, 2021). Normal distribution of the variable in the RS Turipaná, RS Motilonia, RS Carmen de Bolivar and the RS Caribia arranged in violin-plot. **(A)** *NP: number of pods per plant.* **(B)** *NS: average number of seeds per pods.* **(C)** *YLP: yield per plant (g/plant).* **(D)** *SB: seed weight (g).* **(E)** *VB: vegetative biomass (g).*

### All 15,645 SNP Markers were Recovered from an interspecific panel using GBS

DNA extraction at 87 accessions (Table S4) reported a mean DNA concentration (ug/uL) of 3968.80 (IC: 388.02) using Nanodrop^®^, Qubit^®^ mean concentration (ug/uL) of 95, 20 (CI: 8.14), mean A260/280 ratio of 2.13 (CI: 0.01), and mean A260/230 ratio of 2.15 (CI: 0.01). Subsequently, the 87 genetic libraries built for sequencing present: a mean Qubit^®^ concentration (ug/uL) of 16.70 (CI: 2.70), a mean fragment size (bp) of 323.00 (CI: 2.59), and mean TapeStation^®^ quantification of 78.74 nM (CI: 13.00) (Table S5). The electropherograms for each material suggested the distribution of the fragments with defined peaks, and without the presence of contaminants. Only genotype 85 did not have sufficient quality parameters to integrate the process of SNP calling. Following the GATK4 protocol and using the second version of the reference genome of *Phaseolus vulgaris* L. (Schmutz et al., 2014b), a raw matrix of allelic variants was obtained with 1,919,875 sites. After the filters by missing data for loci of 80%, missing data for site of 80%, minimum depth of 3X and MAF (Minimum Alellic Frecuency) of 5%, the final dataset of molecular variants for genetic analysis had the 15,645 SNP markers.

### Genetic Structure and Kinship Relationships Suggested Five Demographic Groups and Substructure Related to Families of Genotypes

The population stratification carried out by the clustering methodologies and ancestry -model approach suggested that all GWAS panel was distributed in five demographic groups (Figure S7-B-C). In the same way, the reconstruction of kinship using VanRaden algorithm suggested 5 groups (Figure S8). However, the GWAS panel was composed by closely related genotypes due to the recurring crosses into the breeding programs. In this sense, the genetic structure used in the GWAS analysis was constructed using a substructure of 12 groups when all genetic groups had at least one genotype with a threshold of greater than 90 percent purity (Figure S7-A). The use of the population structure as covariates related with families of genotypes instead of population structure related with demographic paradigm, more efficiently controlled the rate of false positives, by lowering the p-values of SNPs arranged as a horizontal row, possibly due to the use of covariates in the population stratification.

### A Total of 47 unique Loci in Four Locations with Five Yield Traits Indices were detected

A Total of 47 unique Loci in Four Locations with Five Yield Traits Indices. From the 47 associated SNP markers, 43 were associated with a specific annotated gene in the reference genome of *Phaseolus vulgaris* L storage in *Phytozomne*. The 43 SNPs related with genes, 80 were associated with YLP index, 52 with NS index, 45with SB index, 42 with NP index, and 31 with VB index. Thus, RS Turipaná was the one with the most associated molecular markers (128 SNPs), RS Motilonia and RS Caribia had similar counts of associated SNP markers (60 and 56 respectively), and RS Carmen de Bolivar was the one with the least associated molecular markers (6). The GWAS algorithm BLINK always had lower P-values than the algorithm FarmCPU.

Both GWAS methods recovered similar counts of associated molecular markers (128 SNPs with BLINK and 122 SNPs with FarmCPU) (Table S6), arranged graphically in the circular Manhattan plots (Figure 2–6) with the QQ-plots associated (Figure S9-13)

**Figure 2.**
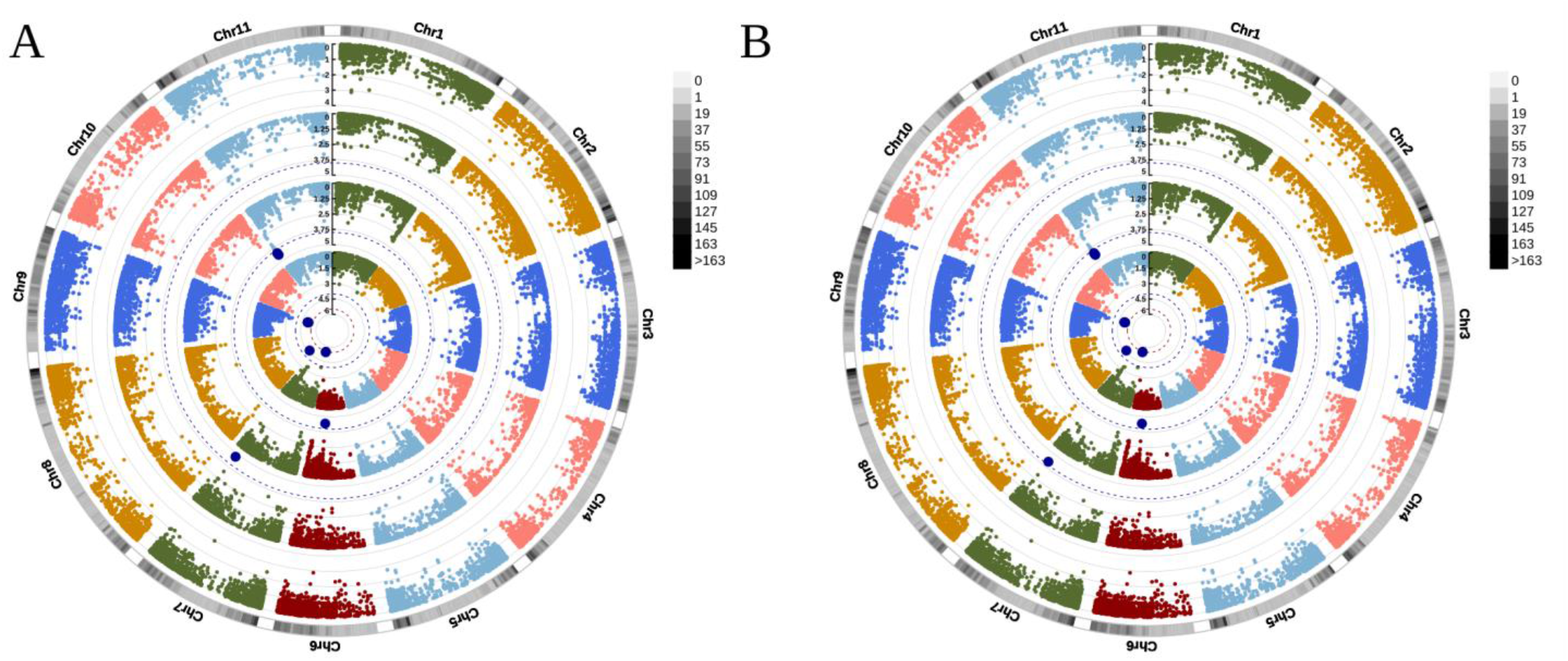
Manhattan plot of multi-environment genome-wide association studies of **pod numbers per plant** using an integrated advanced genotyping panel with common bean (*Phaseolus vulgaris* L.) × tepary bean (*P. acutifolius* A. Gray) interspecific lines in the humid and dry subregions of the Colombian Caribbean. Each ring represents the GWAS results in each locality, from the outer to the inner ring, respectively: RS Turipaná, RS Motilonia, RS Carmen de Bolívar, RS Caribia. In gray the density of genetic markers is observed. Red dots are SNPs that exceeded the Bonferroni threshold (red dotted line, P-value = 3,196 E-6), blue dots are SNPs that exceeded the soft threshold (blue dotted line, P-value = 3,196 E-4). **(A)** *BLINK Algorithm.* **(B)** *FarmCPU Algorithm.*

**Figure 3.**
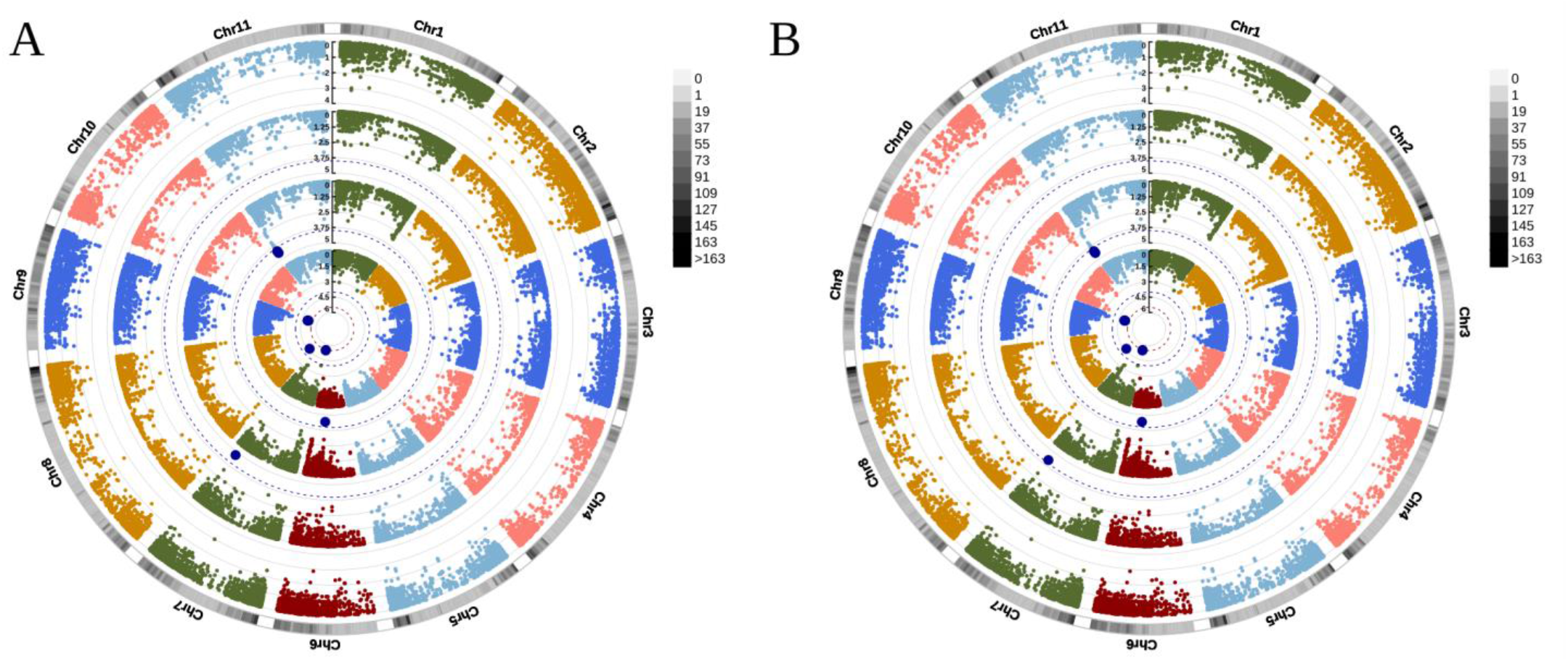
Manhattan plot of multi-environment genome-wide association studies of **average number of seeds per pods** using an integrated advanced genotyping panel with common bean (*Phaseolus vulgaris* L.) × tepary bean (*P. acutifolius* A. Gray) interspecific lines in the humid and dry subregions of the Colombian Caribbean. Each ring represents the GWAS results in each locality, from the outer to the inner ring, respectively: RS Turipaná, RS Motilonia, RS Carmen de Bolívar, RS Caribia. In gray the density of genetic markers is observed. Red dots are SNPs that exceeded the Bonferroni threshold (red dotted line, P-value = 3,196 E-6), blue dots are SNPs that exceeded the soft threshold (blue dotted line, P-value = 3,196 E-4). **(A)** *BLINK Algorithm.* **(B)** *FarmCPU Algorithm.*

**Figure 4.**
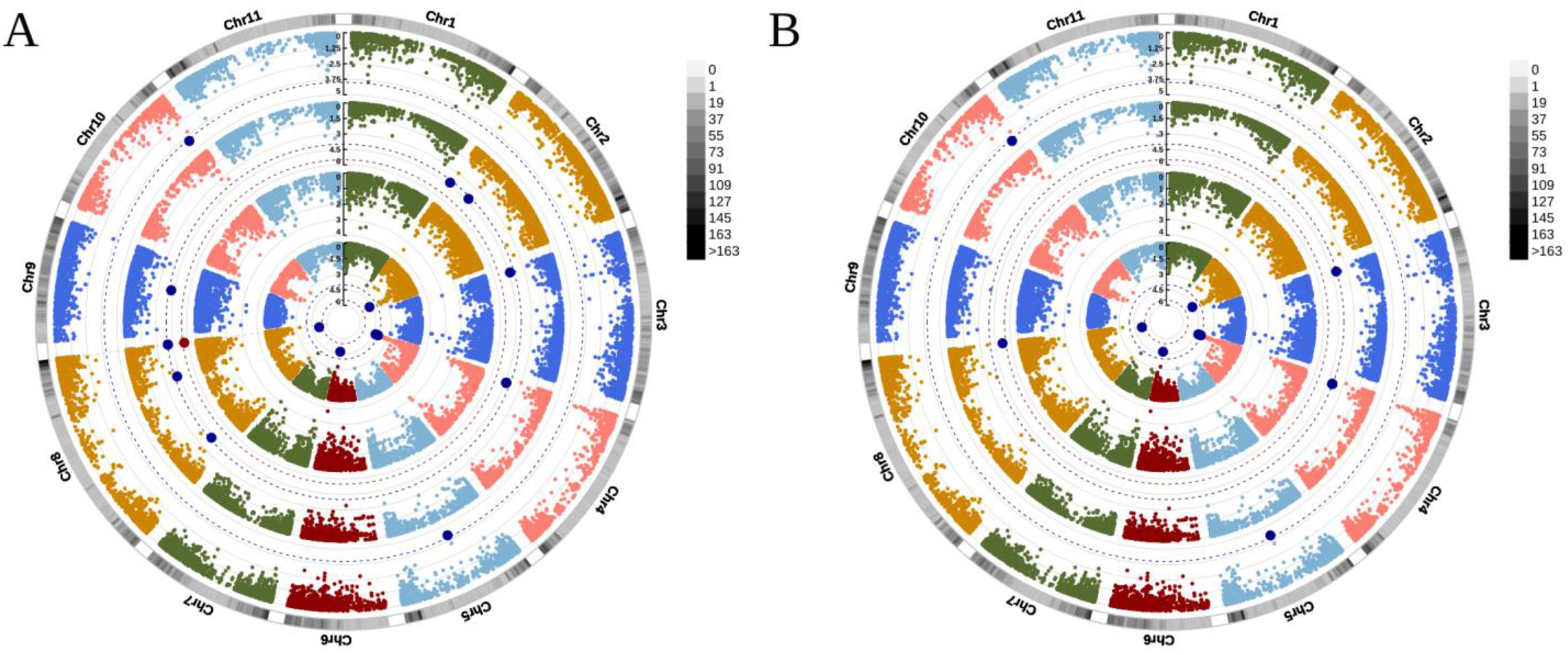
Manhattan plot of multi-environment genome-wide association studies of **yield per plant (g/plant)** using an integrated advanced genotyping panel with common bean (*Phaseolus vulgaris* L.) × tepary bean (*P. acutifolius* A. Gray) interspecific lines in the humid and dry subregions of the Colombian Caribbean. Each ring represents the GWAS results in each locality, from the outer to the inner ring, respectively: RS Turipaná, RS Motilonia, RS Carmen de Bolívar, RS Caribia. In gray the density of genetic markers is observed. Red dots are SNPs that exceeded the Bonferroni threshold (red dotted line, P-value = 3,196 E-6), blue dots are SNPs that exceeded the soft threshold (blue dotted line, P-value = 3,196 E-4). **(A)** *BLINK Algorithm.* **(B)** *FarmCPU Algorithm.*

**Figure 5.**
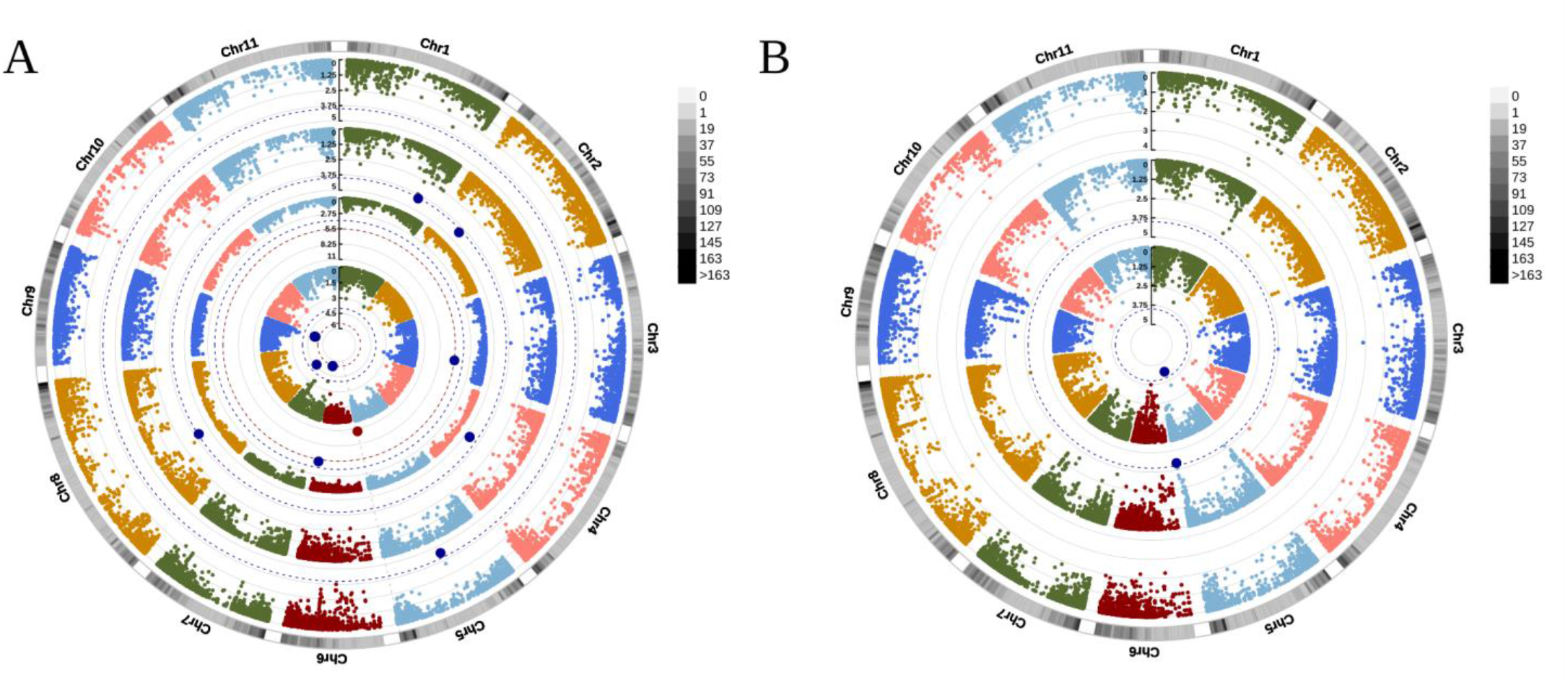
Manhattan plot of multi-environment genome-wide association studies of **seed weight (g)** using an integrated advanced genotyping panel with common bean (*Phaseolus vulgaris* L.) × tepary bean (*P. acutifolius* A. Gray) interspecific lines in the humid and dry subregions of the Colombian Caribbean. Each ring represents the GWAS results in each locality, from the outer to the inner ring, respectively: RS Turipaná, RS Motilonia, RS Carmen de Bolívar, RS Caribia. In gray the density of genetic markers is observed. Red dots are SNPs that exceeded the Bonferroni threshold (red dotted line, P-value = 3,196 E-6), blue dots are SNPs that exceeded the soft threshold (blue dotted line, P-value = 3,196 E-4). **(A)** *BLINK Algorithm.* **(B)** *FarmCPU Algorithm.*

**Figure 6.**
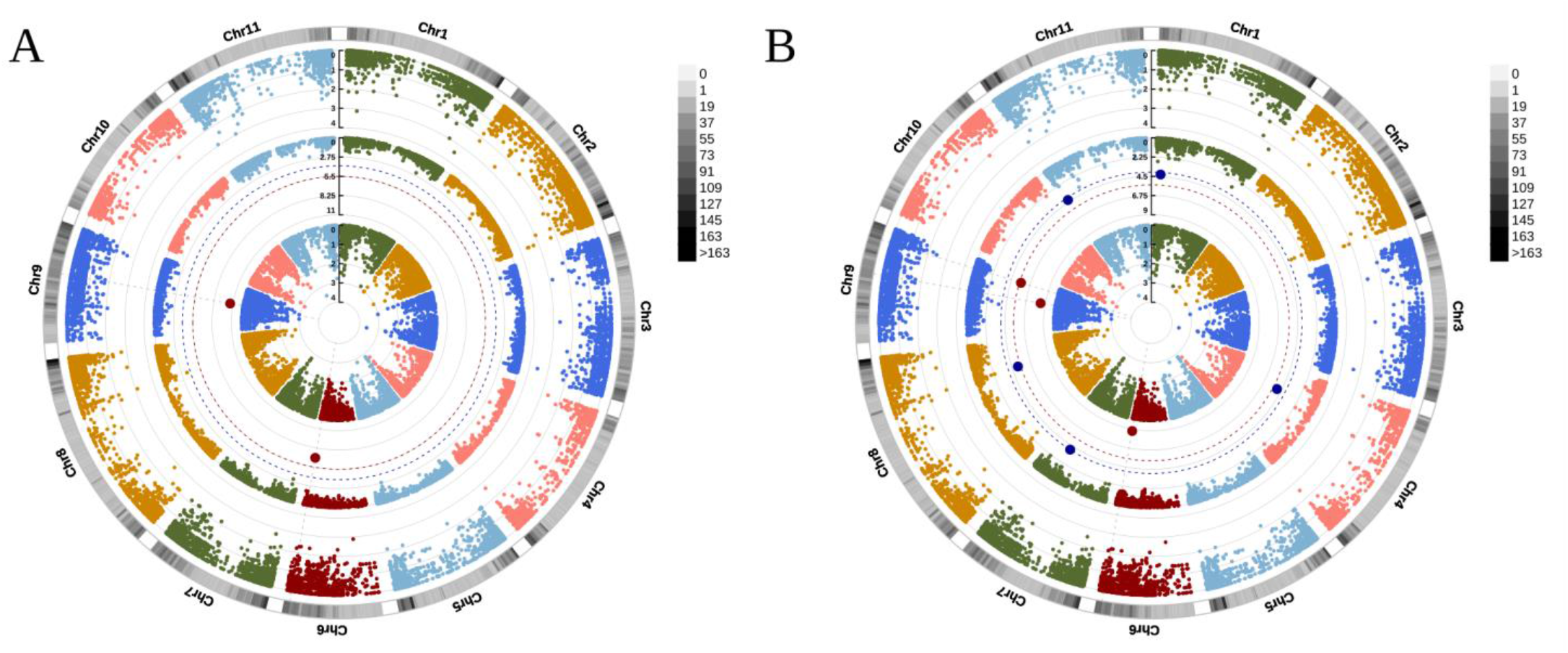
Manhattan plot of multi-environment genome-wide association studies of **pod vegetative biomass (g)** using an integrated advanced genotyping panel with common bean (*Phaseolus vulgaris* L.) × tepary bean (*P. acutifolius* A. Gray) interspecific lines in the humid and dry subregions of the Colombian Caribbean. Each ring represents the GWAS results in each locality, from the outer to the inner ring, respectively: RS Motilonia, RS Carmen de Bolívar, RS Caribia. In gray the density of genetic markers is observed. Red dots are SNPs that exceeded the Bonferroni threshold (red dotted line, P-value = 3,196 E-6), blue dots are SNPs that exceeded the soft threshold (blue dotted line, P-value = 3,196 E-4). **(A)** *BLINK Algorithm.* **(B)** *FarmCPU Algorithm.*

### A 43 Associated Markers Flanked 90 Genes across the All Chromosomes

The chromosomes 11, 9, 8, 7, and 4 had the most count of associated molecular markers with 28, 51, 39, 52, and 35 SNPs respectively. The chromosomes 6, 5, and 2 had 16, 12, and 8 molecular markers respectively. The chromosomes 10, 3, and 1 had the less count of associated molecular markers with 2, 3, and 4 SNPs respectively (Table S6). On the other hand, from the 90 genes, 19 were present in the NP index, 20 in the NS index, 38 in the YLP index, 22 in the SB index, and 27 in the VB index. In the environment context, from the 90 genes, 33 were present in the RS Motilonia, 29 in the RS Caribia, 26 in the RS Turipaná, and 2 in Carmen de Bolivar (Figure 7A).

**Figure 7.**
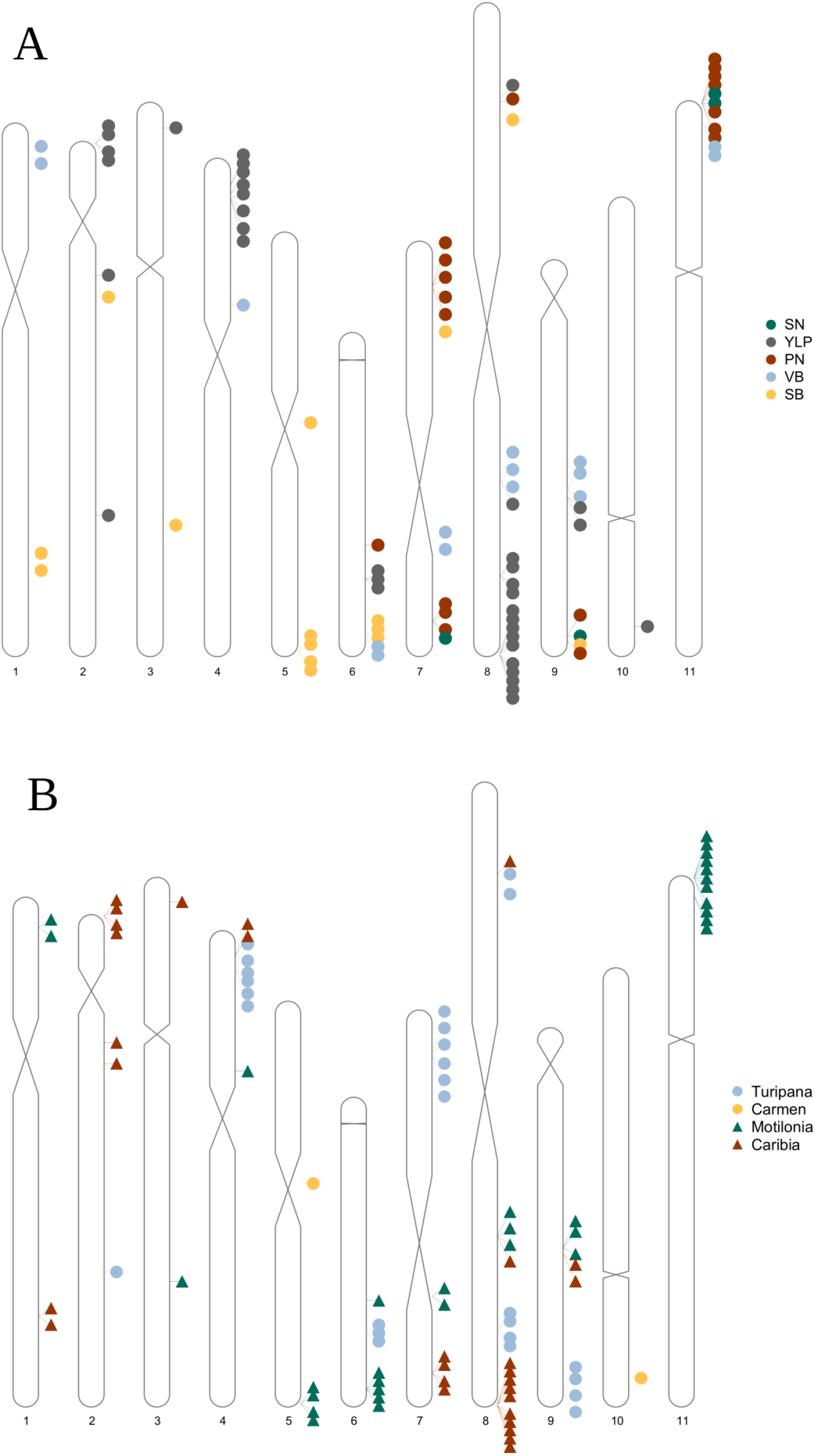
Genetic architecture of the multi-environment GWAS of yield traits in advanced genotypes panel integrated with common bean (*Phaseolus vulgaris* L.) × Tepary Bean (*P. acutifolius* A. Gray) Interspecific Lines at the four localities in the Caribe. **(A)** Idiogram of the five yield traits (*NP: number of pods per plant, NS: average number of seeds per pods, YLP: yield per plant, SB: seed weight, VB: vegetative biomass*) **(B)** Idiogram of the four localities (RS Turipaná, RS Motilonia, RS Carmen de Bolívar, and RS Caribia).

### The associated genes are related to the response mechanisms of drought stress to level morphological, physiological, metabolic, signal transduction, and fatty acid and phospholipid metabolism

The 90 Genes flanked by 43 associated loci suggested several response mechanisms of drought stress. In dry subregions the response mechanism to drought stress was signal transduction (*i.e. Ethylene Response Factor ERF),* photosynthesis (*i.e. LHCA3, chloroplast stem-loop binding protein* and *chloroplastic mate efflux family protein 3*), and drought-induced proteins (*i.e. Late embryogenesis abundant).* On the other hand, in humid subregions, the genetic base of drought tolerance was composed by morphology and physiology (*i.e. wax inducer1/shine1),* photosynthetic capacity (*i.e. Plastocyanin-like domain),* drought-Induced proteins (*i.e. MYB),* reactive Oxygen metabolism (*i.e. peroxidase proteins),* and fatty acid and phospholipid metabolism.

### Pathways Enriched for Genes Associated in the NP, NS, SB, and VB Indices: Superpathway of Fatty Acid Biosynthesis II, Stearate Biosynthesis II, and Palmitate Biosynthesis II

Pathway enrichment analysis suggested 28 metabolic pathways enriched for genes using the platform Phytozome. Three pathways were found in the NP, NS, SB, and VB Indices related to The Fatty Acid and Phospholipid Metabolism: *Palmitate Biosynthesis II, Stearate Biosynthesis II,* and *Superpathway of Fatty Acid Biosynthesis II.* The NP index and NP index captured the same metabolic pathways: *cis-vaccenate biosynthesis, fatty acid elongation*, *gondoate biosynthesis* (anaerobic), *octanoyl-[acyl-carrier protein] biosynthesis*, *palmitate biosynthesis II, palmitoleate biosynthesis I, stearate biosynthesis II,* and the *superpathway of fatty acid biosynthesis II.* The NS index captured two pathways more than NP index: *adenosine ribonucleotides de novo biosynthesis, and the superpathway of adenosine nucleotides de novo biosynthesis I.* Also, the SB index and VB index captured the same metabolic pathways: *cutin biosynthesis, fatty acid & beta-oxidation II* (peroxisome), *long-chain fatty acid activation, oleate biosynthesis I*, *palmitate biosynthesis II*, *phosphatidylcholine acyl editing*, *phytol degradation*, *sporopollenin precursors biosynthesis, stearate biosynthesis II, suberin monomers biosynthesis, superpathway of fatty acid biosynthesis II,* and the *wax esters biosynthesis II.* The SB index captured one pathway more than VB index: *superpathway of glyoxylate cycle and fatty acid degradation*. On the other hand, the YLP index captured several pathways: *baicalein degradation* (hydrogen peroxide detoxification), *justicidin B biosynthesis*, *L-glutamine biosynthesis III*, *luteolin triglucuronide degradation*, *matairesinol biosynthesis*, *phenylpropanoid biosynthesis*, *initial reactions, superpathway of scopolin and esculin biosynthesis,* and the *TCA cycle II.* (Table 1).

**Table 1.**
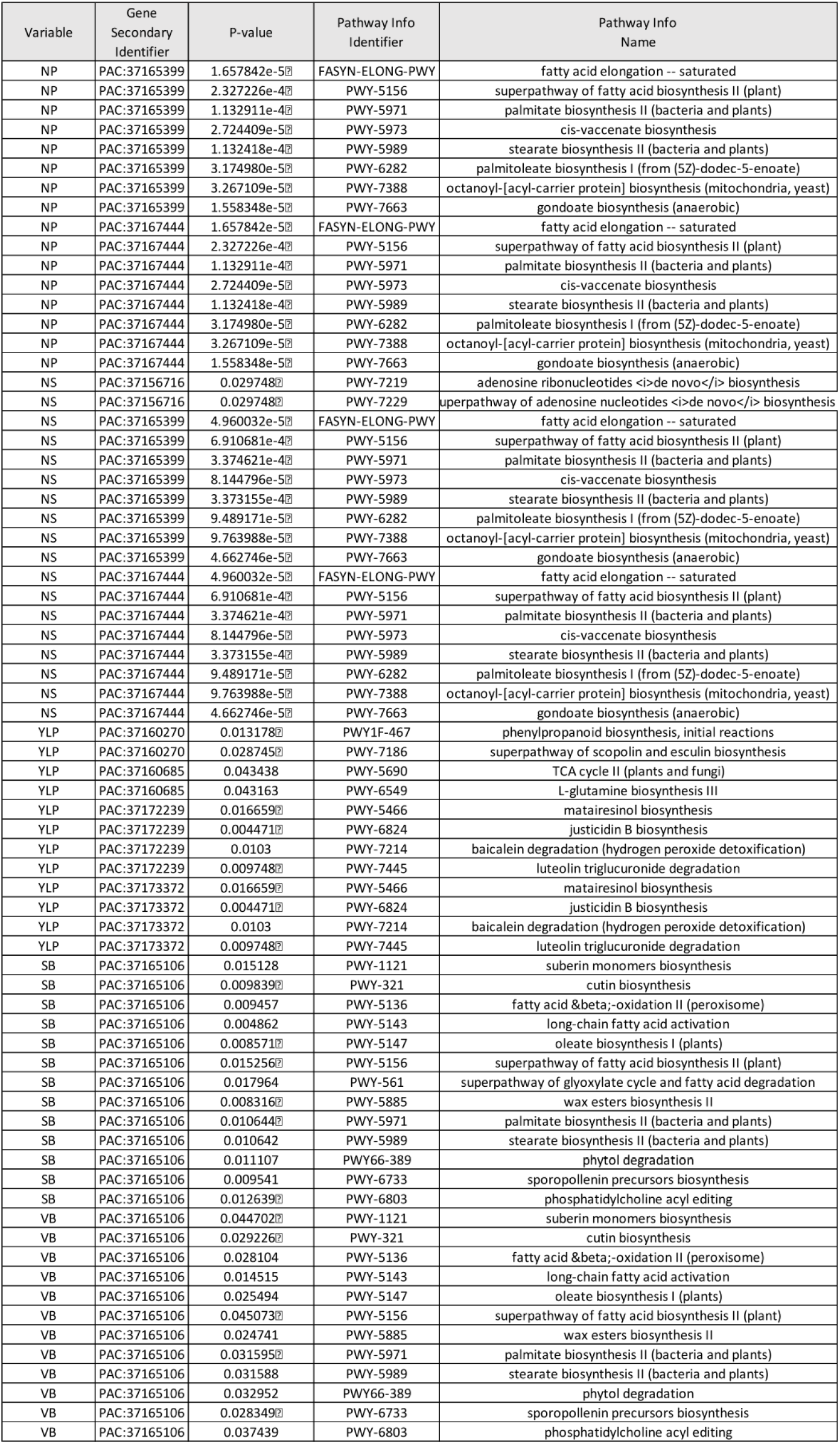
Pathways enriched analysis for each yield traits indices across the environments in the Caribe. The software used was the PhytoMine tool in the Phytozome v. 13 platform. The p-value was calculated using the Hypergeometric distribution.

## DISCUSSION

The understanding of adaptive genetic variation to drought and high temperature stress in the Caribbean subregions, which has arid and semi-arid ecosystems, have been of great interest in the development of promising breeding programs to power the food security in the region. However, the advanced panels with interspecific genotypes and modern GWAS approaches have given importance to the selection of SNP markers. The five indices related to yield traits (NP, NS, YLP, SB, and VB) and GWAS models (BLINK and FarmCPU) that generated significant results captured complementary components of the genetic architecture of drought tolerance. We found a total of 43 loci associated to 90 genes related to biological processes of the drought tolerance response in plants under humid and dry regions of the Caribe. Also, from the 90 genes, we captured several enrichment pathways for genes centered in the fatty acid biosynthesis, and this relationship has been associated to abiotic stress response, such as drought and heat stress (Asakura et al., 2021). The loci captured in this work could be potential candidates to powered breeding programs using molecular marker-guided selection, genome editing, genomic selection, *inter alia*.

### Morphology and Physiology as Response Mechanism to Drought and high temperature Stress

The response mechanism of plants to drought stress has been studied on plant morphology and physiology (Yang et al., 2021). Therefore, in drought stress, the internal structure and physical property of plants change and adapt. For instead, the cuticle is a kind of lipid membrane, which can reduce the loss of water to the atmosphere, acting as a barrier for plant water evaporation. Their cuticle is composed by several biomolecules as cutin and soluble waxes. In a wide range of plants, the altering the cutin and suberin includes increased sensitivity to biotic and abiotic stresses, defects in development and growth, and changed morphology, permeability, and seed dormancy (Pollard et al., 2008), being consistent with the enrichment pathway reported *suberin monomers biosynthesis* for vegetative traits (SB and VB). Also, tea leaves improve drought resistance through increasing wax coverage, cuticle thickness, and osmiophilicity (Patwari et al., 2019; Chen et al., 2020), when we reported the enrichment pathway *wax esters biosynthesis II.* Additionality, we found that the SNP markers *S04 3848215, S04_3848227,* and *S04_3848215* were associated with the gen *Phvul.004G031900,* a transcription factor (*wax inducer1/shine1*) of the Ethylene Response Factor (ERF) family, that has recently been shown to induce the production of epidermal waxes when overexpressed in *Arabidopsis* plants (Kannangara et al., 2007) and *Brassica napus* (Liu et al., 2019). The previous observation that chitinase genes are involved in both leaf development and senescence (Quirino et al., 2000) points to a gene in this cluster as strong candidate gene for the whole plant response under heat stress.

### Metabolic Pathways as Response Mechanism to Drought and high temperature Stress

The drought and high temperature stress affect the physiological and biochemical characteristics as: Photosynthetic Capacity, Drought-Induced Proteins, and Reactive Oxygen Metabolism (Yang et al., 2021). Photosynthesis is one of the main processes affected by water stress. We found in all research stations genes related to photosynthesis metabolism as: *Plastocyanin-like domain* (Phvul.007G055000) in RS Turipaná and the genes *LHCA3* (Phvul.008G289700), *chloroplast stem-loop binding protein* (Phvul.007G172100) and *chloroplastic mate efflux family protein 3* (Phvul.008G291400) in the dry RS. Also, we obtained the pathway enrichment in genes: *phytol degradation.* On the other hand, drought-induced proteins play a protective role in plant adaptation to stress and can improve plant drought tolerance. These drought-induced proteins are divided in functional proteins and regulatory proteins. We found a relevant protective protein named *Late embryogenesis abundant* (LEA) (Chen et al., 2021) (Phvul.011G001700) only in the RS Motilonia using the SNP marker S11_140976, in other legume of importance such as *Vigna sp.* has been reported up regulation of some genes grouped like LEA in heat stress tolerant genotypes (Singh et al., 2022). Therefore, we found the regulatory protein *MYB* in RS Turipaná (Phvul.008G205000) and Caribia (Phvul.002G088900), particularly a tranbscription factor in the *biosynthesis of phenylpropanoids*, an enrichment pathway that we found too (Deng and Lu, 2017; Yang et al., 2021). In the context of Reactive Oxygen Metabolism, we found peroxidase protein related to removal of H2O2 (Gill and Tuteja, 2010) (Phvul.006G130000) in the RS Turipaná, and the enrichment pathway *baicalein degradation* (hydrogen peroxide detoxification).

### Signal Transduction as Response Mechanism to Drought and high temperature Stress

The plants under drought stress implement signal transduction (Asakura et al., 2021). We found the gene of ERF (Phvul.005G180700) in the RS Motilonia. The Ethylene Response Factor (ERF) play an important role in plant stress resistance and previous studies (Kannangara et al., 2007; Cheng et al., 2013) have shown that they can participate in the process of drought stress resistance in plants through different pathways by affecting the synthesis of plant hormones as jasmonic acid and ethylene (Yang et al., 2021). In the same way, these hormones participate in signaling processes between plants, in response to other types of stresses, which are also considered of great importance in the face of projections of the negative impact of climate change, such as biotic stresses.

### Fatty Acid and Phospholipid Metabolism as Response Mechanism to Drought and high temperature Stress

For decades, the alteration in *fatty acid* and *phospholipid metabolism* have been reported in legumes under drought stress (Dornbos and Mullen, 1992). Recently, (Asakura et al., 2021) explored the regulation of *fatty acid* and *phospholipid metabolism* in tomato fruit under drought stress using transcriptomic and metabolomic analyses. The main result of these authors was that *fatty acid* and *phospholipid metabolism* are a component in several metabolic pathways of drought tolerance as: Cell wall, Ethylene and Jasmonate biosynthesis (Kazan, 2015). We found several enrichment pathways (RS Turipaná and RS Caribia) that were the same obtained by (Asakura et al., 2021) as the *superpathway of fatty acid biosynthesis II, palmitate biosynthesis II,* and *stearate biosynthesis II,* captured simultaneously by the yield traits indices NP, NS, YLP, SB, and VB. These pathway networks could be useful like a bioindicator with potential to health impact in interspecific lines of bean with Tepary and common bean like parentals, under drought conditions. Therefore, the results from structure analysis confirmed the two base panels represent distinct populations and are appropriate for studies designed to investigate the genetic factors controlling important agronomic traits within each gene pool. This SNP data set will allow researchers to determine whether traits are controlled by genetic factors shared by both gene pools or whether gene pool specific factors are controlling important traits (Oladzad et al., 2019).

### Caribbean Subregions Suggest Differences in Genetic Base of Drought and high temperature Tolerance

We observed that in the Caribbean subregions, the genetic base of drought and high temperature tolerance could be different. For example, in dry subregions the response mechanism to drought stress was signal transduction, photosynthesis and drought-induced proteins. On the other hand, in humid subregions, the genetic base of drought tolerance was composed by morphology and physiology, photosynthetic capacity, drought-Induced proteins, reactive Oxygen metabolism, and fatty acid and phospholipid metabolism. However, in all locations of Caribe, the response mechanism common was related to the photosynthesis pathway, but different in the proteins associated. Finally, we recommend the use of advanced panels with interspecific genotypes and modern GWAS approaches in order to reveal the adaptive genetic variation of drought and high temperature stress across the environments as well as Caribbean subregions. Finally, the use of enrichment analysis pathways could help to analyze wide panel of associated loci and they can be used as selection tools for traits important for high crop productivity and it can be used to search for genetic factors and polymorphisms that would be useful for improvement in their breeding programs.

## PERSPECTIVES

This study demonstrates that the implementation an advanced panels with interspecific genotypes and modern GWAS models under different environments with carefully yield traits indices improves the reconstruction of the genetic basis of adaptation to heat and drought stress. New studies across a variety of novel interspecific genotypes subjected to different stresses will benefit by using GWAS models within a well thought environmental design in order to capture better sources of genetic adaptation. We are looking forward to seeing more studies that follow these lines within the oncoming years.

Instead, the genes identified as candidates for drought tolerance have the potential to be used in plant breeding programs after validation by means of strategies such as gene expression studies and Whole Genome re-Sequencing (WGS) (Barbulescu, D. M., 2018). Thus, the validated candidate genes could be integrated into molecular editing strategies as CRISPR/Cas9 (Lang-Mladek et al., 2010; Pecinka et al., 2010; LeBlanc et al., 2018) and Genomic Selection (Assefa et al., 2019).

As part of a larger project, promissory accessions identified in this work will be integrated in the coming stages of KoLFACI breeding project for food security in the Colombian Caribbean. Also, as part of this project, the associations SNPs that reconstruct the genetic base will be used as improvement in the Genomic Prediction modeling powered by machine learning approaches to integrate as Genomic Selection process in coming stages.

## AUTHOR CONTRIBUTIONS

**Conceptualization:** F.L.-H., A.J.C., E.B.-E., R.I.L.-P., C.C.C.-C., D.F.V.-M., and A.P.T.-R. **Methodology:** F.L.-H., E.B.-E., R.I.L.-P., C.C.C.-C., A.J.C. and A.P.T.-R. designed sampling and experiments with assistance from S. Beebe. **Data Collection:** R.I.L.-P. and C.C.C.-C. **Data Curation:** F.L.-H. **Bioinformatic & Statistic Scripts:** F.L.-H. **Visualization:** F.L.-H. **Supervision:** A.P.T.-R., D.F.V.-M., and A.J.C. **Writing:** F.L.-H. wrote the manuscript with contributions from A.J.C., E.B.-E., R.I.L.-P., C.C.C.-C., D.F.V.-M., and A.P.T.-R. All authors have read and agreed to the published version of the manuscript.

## ACKNOWLEDGMENTS

The authors express their acknowledgments to the Korea-Latin America Food and Agriculture Cooperation Initiative (KoLFACI) for funding, the Alliance Bioversity–CIAT (International Center for Tropical Agriculture) for providing interspecific congruity-backcross lines between common (P. vulgaris) and Tepary (P. acutifolius) beans, the Colombian Agricultural Research Corporation (AGROSAVIA) for technical assistance, and Ministerio de Agricultura y Desarrollo Rural de Colombia (MADR) for administrative support. Authors also thank S. Beebe and V. Mayor for providing seed material, and the genealogies of the genotypes. F.L.-H. acknowledges insightful discussions around abiotic stress in common bean with M.W. Blair, which took place during the summer of 2019 in Rionegro (Antioquia, Colombia) as part of the funding provided by the Fulbright’s U.S. Specialist Program. F.L. -H. recognizes AGROSAVIA’s Department for Research Capacity Building for subsidizing his internship in 2018, which enabled start exploring G × E models for heat stress in common bean. Authors deeply acknowledge to discussions around drought stress in common bean with M. O. Urban, S. Cruz, J.S. Aparicio, S. Beebe, J. Soto, J. Ricaurte, which took place during the 2022 in Palmira (Valle del Cauca, Colombia). Also, we thank to S. Barrera for inlight the understanding of interspecific genotypes. F.L.-H. acknowledges to J. Berdugo, J. De Vega, K. Denning-James, and CABANA workshop 2021 for discussion around bioinformatic and statistical approaches.

## SUPPLEMENTAL MATERIAL

**Table S1**. Genealogy of 87 genotypes, composed of 67 interspecific lines between the common bean (*P. vulgaris*) and Tepary bean (*P. acutifolius*), and 19 advanced genotypes bred to high temperature and drought conditions by the bean program of the Alliance Bioversity –CIAT. The genotype G40001 (*P. acutifolius*) as control. The panel of genotypes was evaluated for the first time at four localities in the humid and dry Colombian Caribbean subregions (Burbano-Erazo et al., 2021).

**Table S2**. Research Centers’s description and soil fertility scores in coastal Colombia

**Table S3**. The Shapiro-Wilk test output to corroborated the normality behavior in all localities in all yield index using the R-package *nortest* (Gross and Ligges, 2015)

**Table S4**. DNA extraction at 87 accessions

**Table S5.** Genetic libraries built at 87 accessions for sequencing

**Table S6**. Multi-environment GWAS of yield traits indices in advanced genotypes panel integrated with common bean (*Phaseolus vulgaris* l.) × tepary bean (*p. acutifolius* a. gray) interspecific lines at the humid and dry Colombian Caribbean subregions.

**Figure S1**. The Global daily (1km) land surface precipitation based on cloud cover-informed downscaling (Karger et al., 2021), the time series of daily precipitation (kg/(må(2) day)) was extracted for the same months from cultivation to harvest from the historical data of year 2003 to year 2016. **(A)** RS Motilonia **(B)** RS Caribia **(C)** RS Carmen de Bolivar **(D)** RS Turipaná

**Figure S2**. *Number of pods per plant* for the GWAS panel through the four locations of the Colombian Caribbean (A) RS Turipaná (B) RS Motilonia (C) RS Carmen de Bolívar (D) RS Caribia

**Figure S3**. *Number of seeds per pod* for the GWAS panel through the four locations of the Colombian Caribbean (A) RS Turipaná (B) RS Motilonia (C) RS Carmen de Bolívar (D) RS Caribia

**Figure S4**. *Yield per plant (g/plant) for* the GWAS panel through the four localities of the Colombian Caribbean (A) RS Turipaná (B) RS Motilonia (C) RS Carmen de Bolívar (D) RS Caribia

**Figure S5**. *Seed weight (g*) for the GWAS panel through the four locations of the Colombian Caribbean (A) RS Turipaná (B) RS Motilonia (C) RS Carmen de Bolívar (D) RS Caribia

**Figure S6**. *Vegetative biomass (g*) for the GWAS panel through the four localities of the Colombian Caribbean (A) RS Motilonia (B) RS Carmen de Bolívar (C) RS Caribia

**Figure S7.** Population stratification. **(A)** non-negative matrix factorization algorithm (*snmf* function) in the LEA R-package, an improvement to admixture analysis (Frichot and François, 2015). **(B)** Cross-Entropy optimization with 10,000 repetitions of *snmf* function. **(C)** Molecular principal component analysis (hereinafter referred to as PC) carried out by optCluster R-package (Sekula et al., 2017).

**Figure S7.** Heat map of kinship matrix estimated with the VanRaden algorithm across the 15,645 SNP markers.

**Figure S9**. QQ-plot of the multi-environment genome-wide association studies of **pod numbers per plant** using an integrated advanced genotyping panel with common bean (*Phaseolus vulgaris* L.) × tepary bean (*P. acutifolius* A. Gray) interspecific lines in the humid and dry subregions of the Colombian Caribbean. Each color represents the GWAS results in each locality: RS Turipaná, RS Motilonia, RS Carmen de Bolívar, RS Caribia.

**Figure S10**. QQ-plot of multi-environment genome-wide association studies of **number of seeds per pod** using an integrated advanced genotyping panel with interspecific lines of common bean (*Phaseolus vulgaris* L.) × tepary bean (*P. acutifolius* A. Gray) in the humid and dry subregions of the Colombian Caribbean. Each color represents the GWAS results in each locality: RS Turipaná, RS Motilonia, RS Carmen de Bolívar, RS Caribia.

**Figure S11**. QQ-plot of multi-environment genome-wide association studies of **yield per plant (g/plant)** using an advanced genotyping panel integrated with interspecific lines of common bean (*Phaseolus vulgaris* L.) × tepary bean (*P. acutifolius* A Gray) in the humid and dry subregions of the Colombian Caribbean. Each color represents the GWAS results in each locality: RS Turipaná, RS Motilonia, RS Carmen de Bolívar, RS Caribia.

**Figure S12**. QQ-plot of multi-environment genome-wide association studies of **seed weight (g)** using an integrated advanced genotyping panel with common bean (*Phaseolus vulgaris* L.) × tepary bean (*P. acutifolius* A. Gray) interspecific lines) in the humid and dry subregions of the Colombian Caribbean. Each color represents the GWAS results in each locality: RS Turipaná, RS Motilonia, RS Carmen de Bolívar, RS Caribia.

**Figure S13**. QQ-plot of multi-environment genome-wide association studies of **vegetative biomass (g)** using an integrated advanced genotyping panel with interspecific lines of common bean (*Phaseolus vulgaris* L.) × tepary bean (*P. acutifolius* A. Gray) in the humid and dry subregions of the Colombian Caribbean. Each color represents the GWAS results in each locality: RS Motilonia, RS Carmen de Bolívar, RS Caribia.

