## Supplementary figures and images for "Multi-environment Genome Wide Association Studies of Yield Traits in Common Bean (*Phaseolus vulgaris* L.) × Tepary Bean (*P. acutifolius* A. Gray) Interspecific Advanced Lines at the Humid and Dry Colombian Caribbean Subregions"

### Supplemental Figure 1

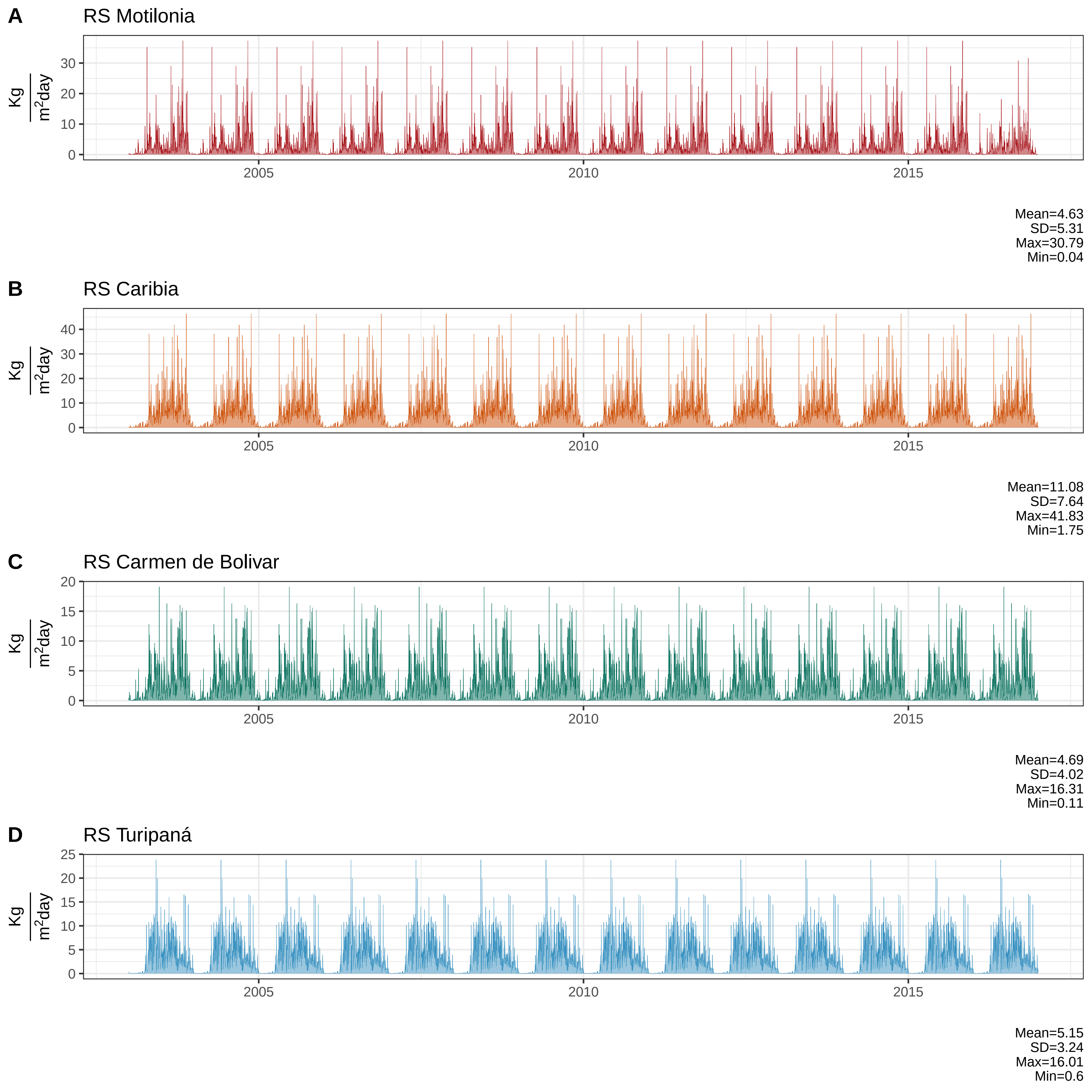

### Supplemental Figure 2

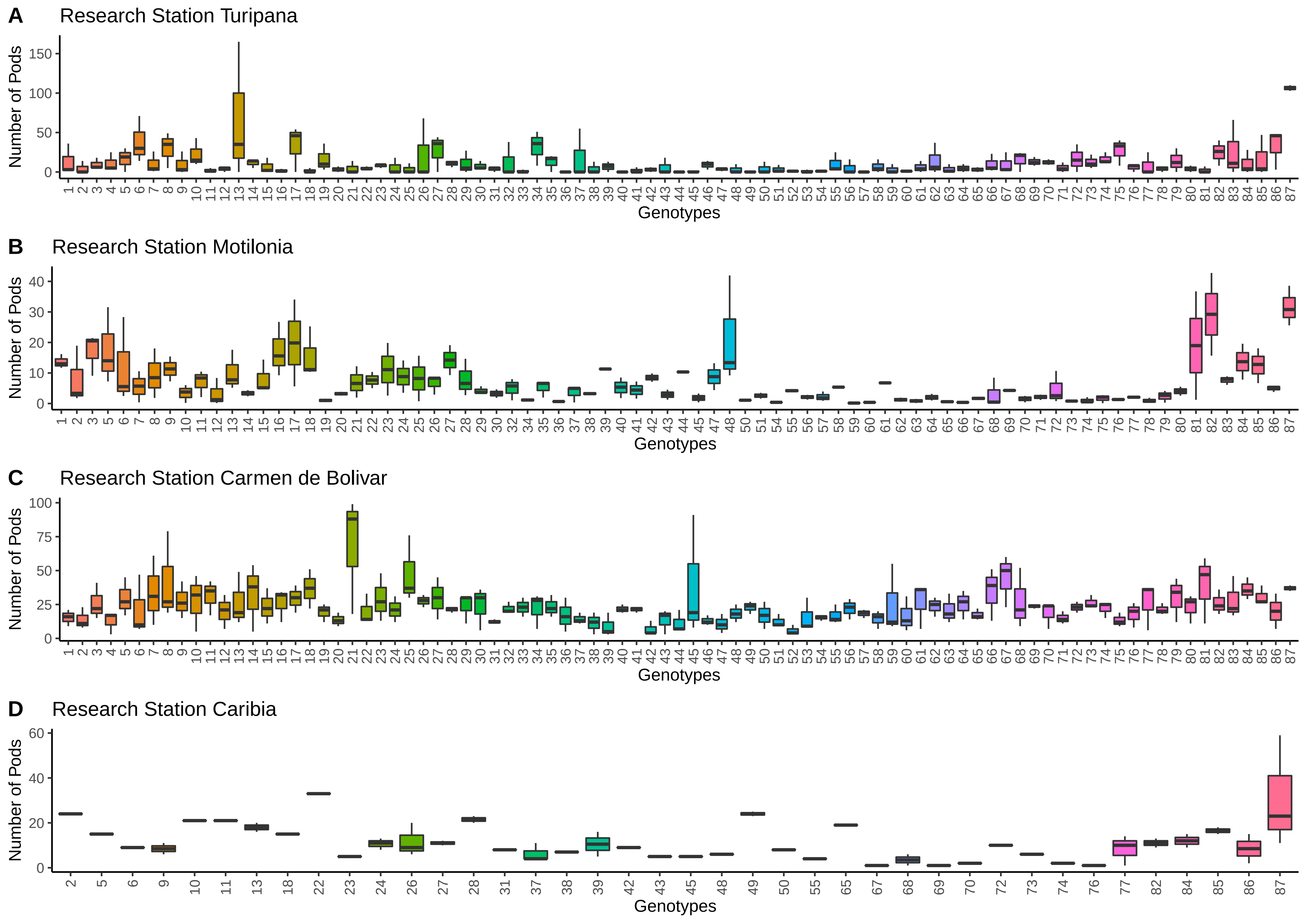

### Supplemental Figure 3

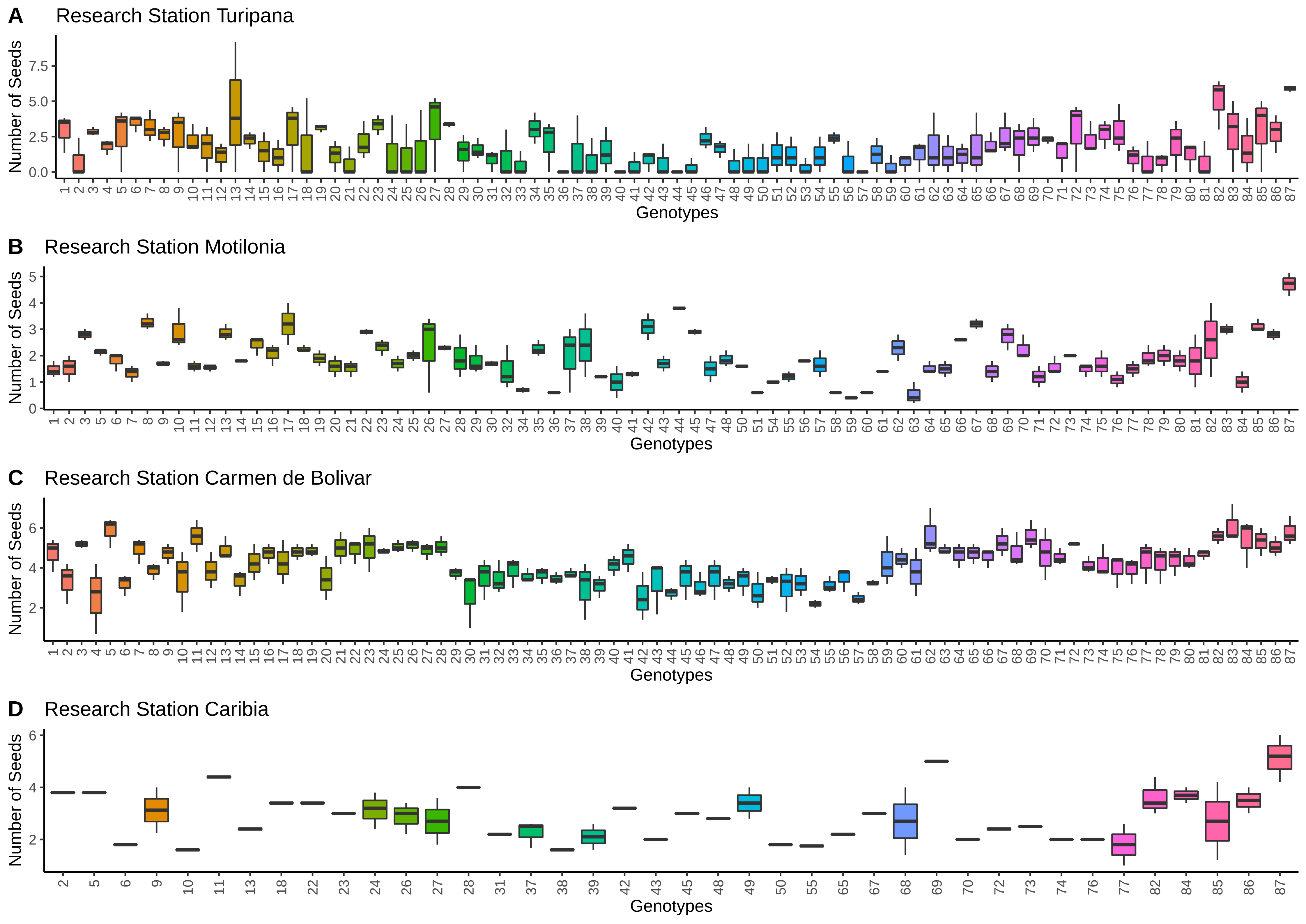

### Supplemental Figure 4

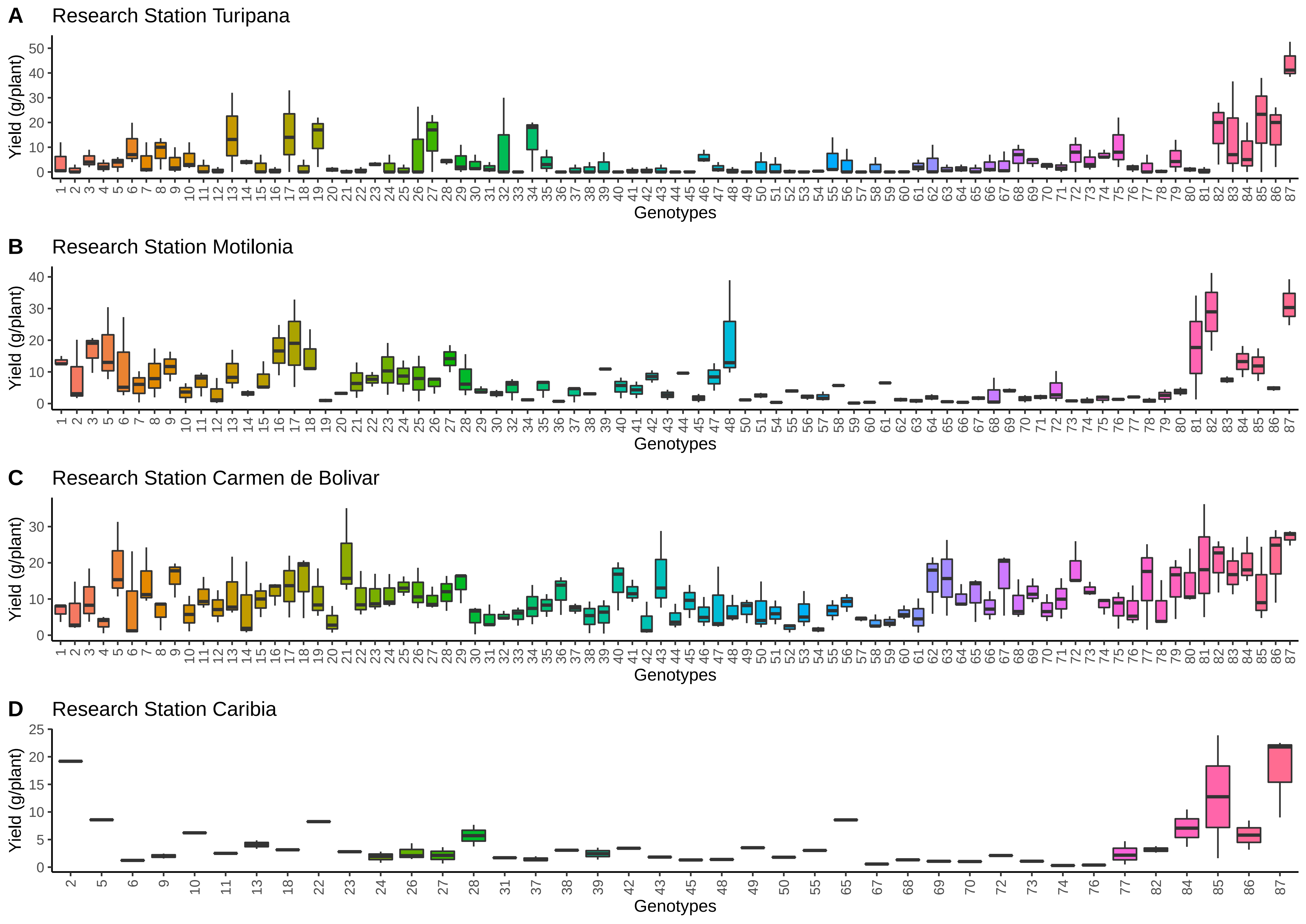

### Supplemental Figure 5

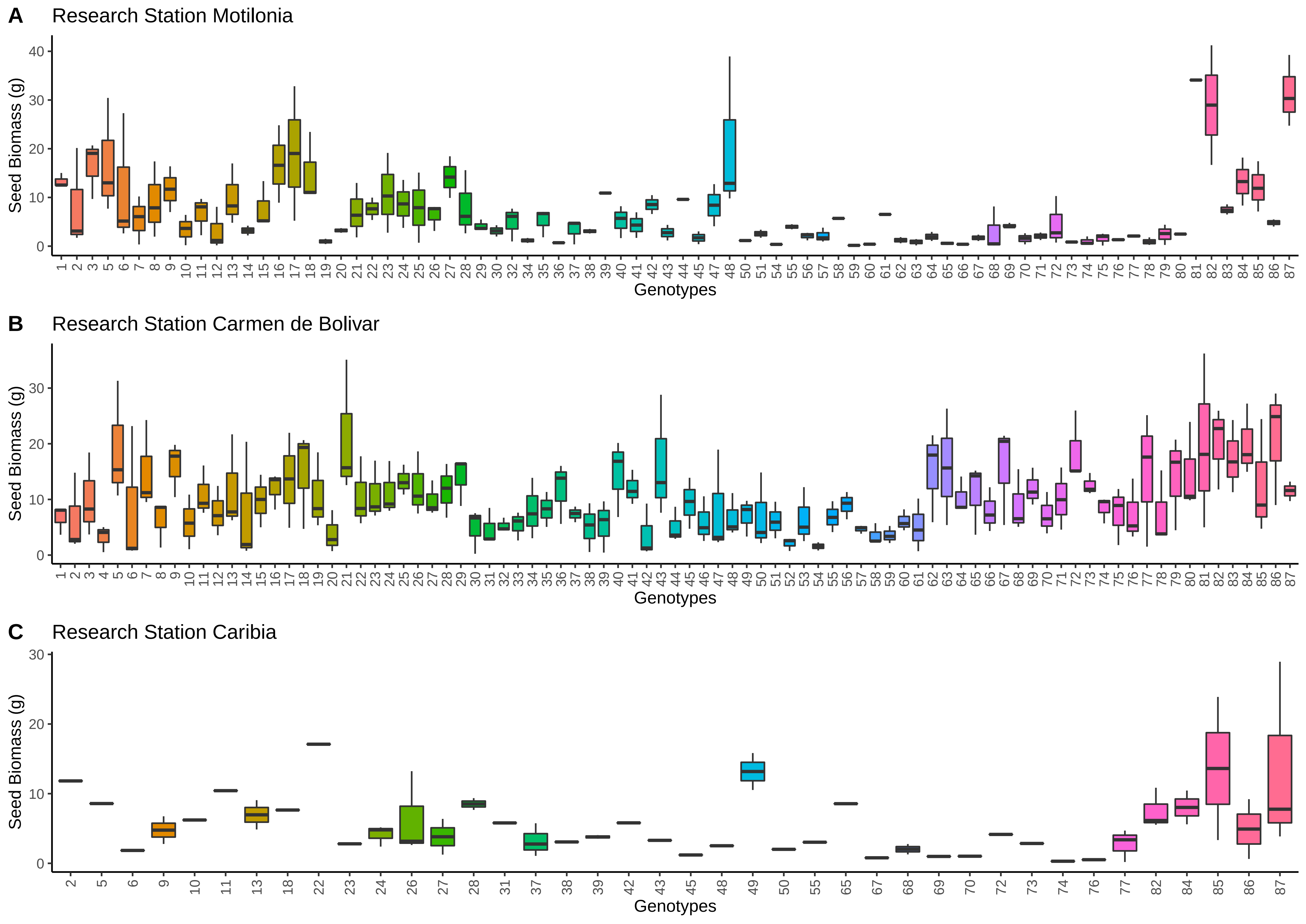

### Supplemental Figure 6

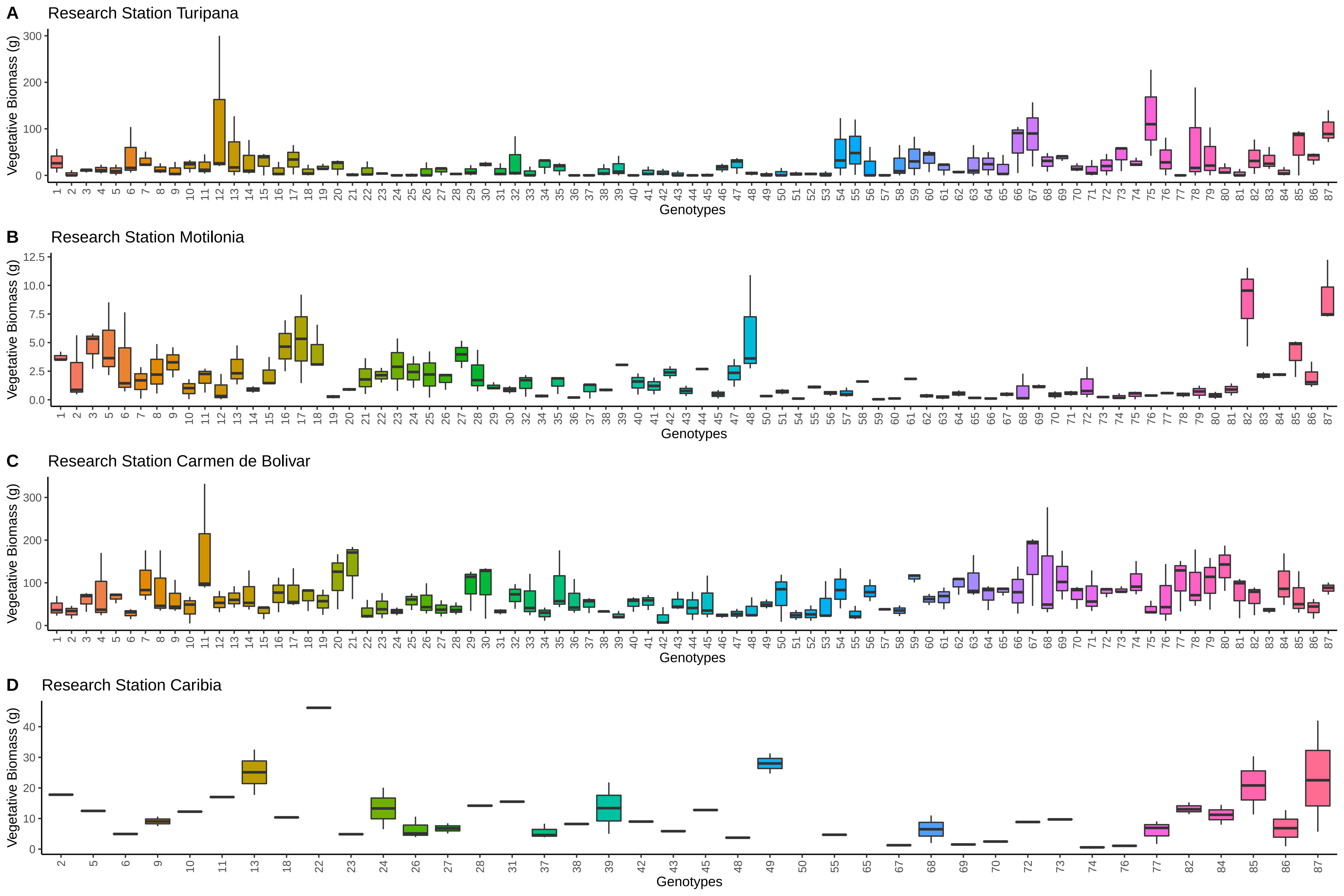

### Supplemental Figure 7

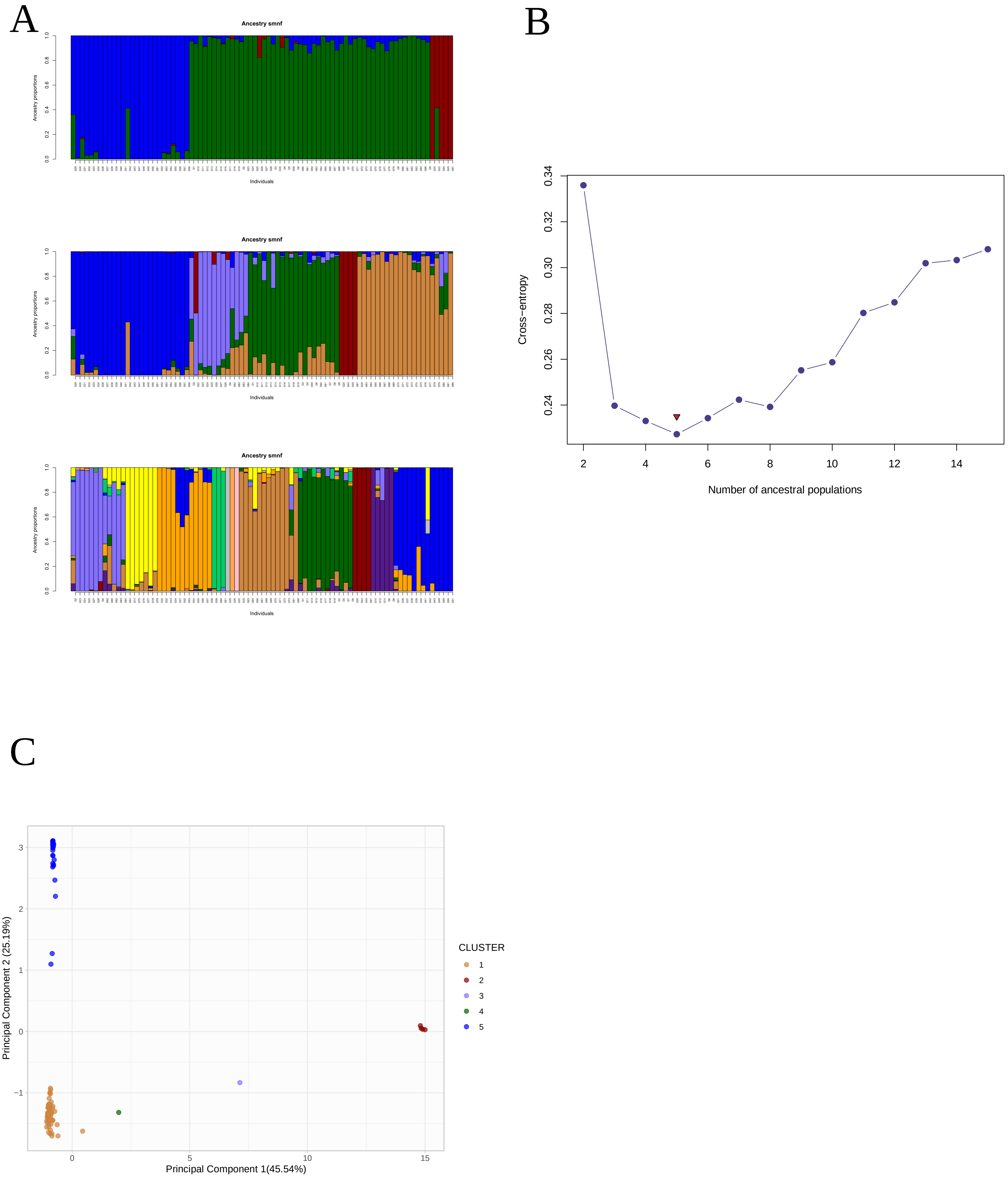

### Supplemental Figure 8

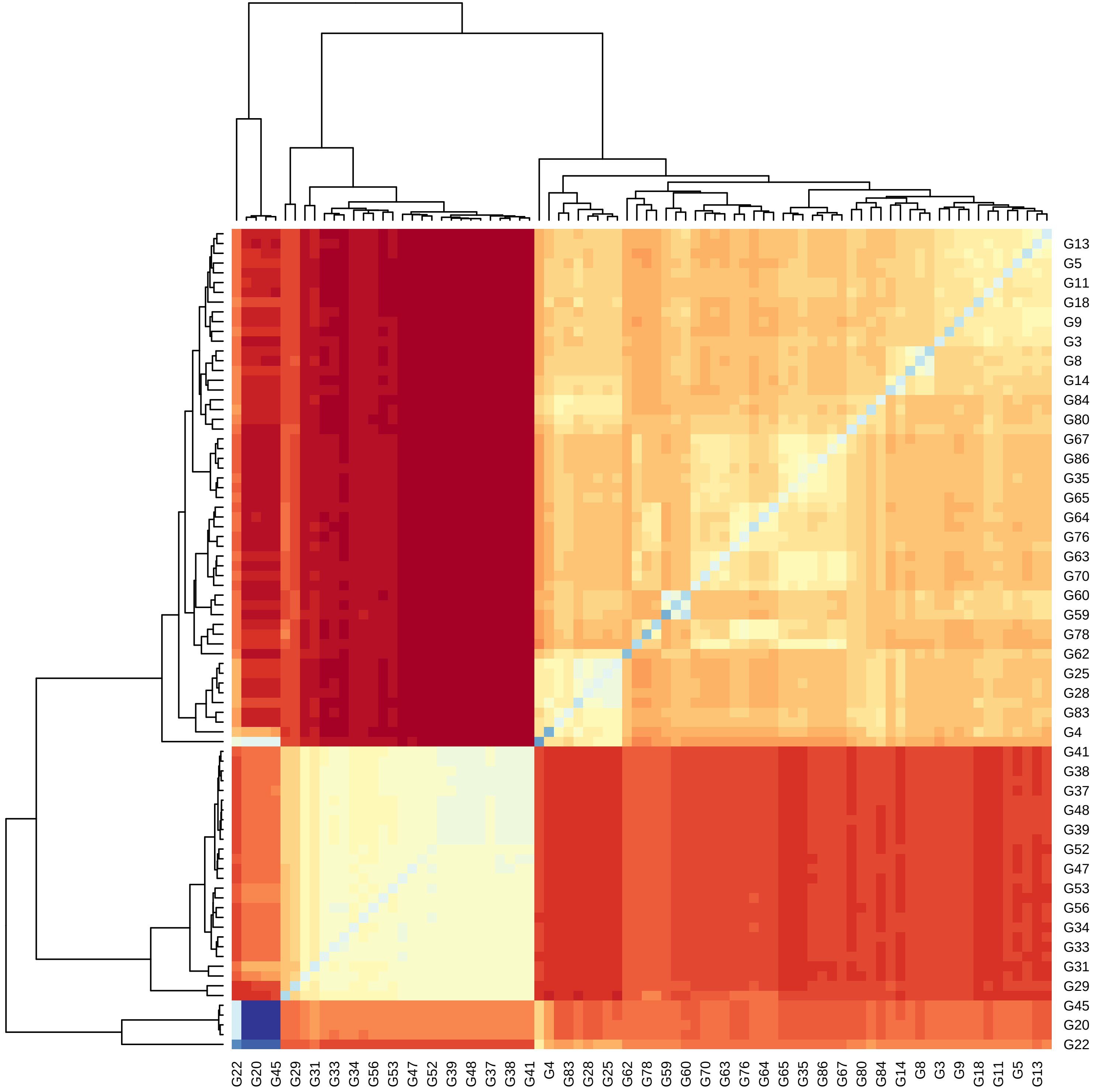

### Supplemental Figure 9

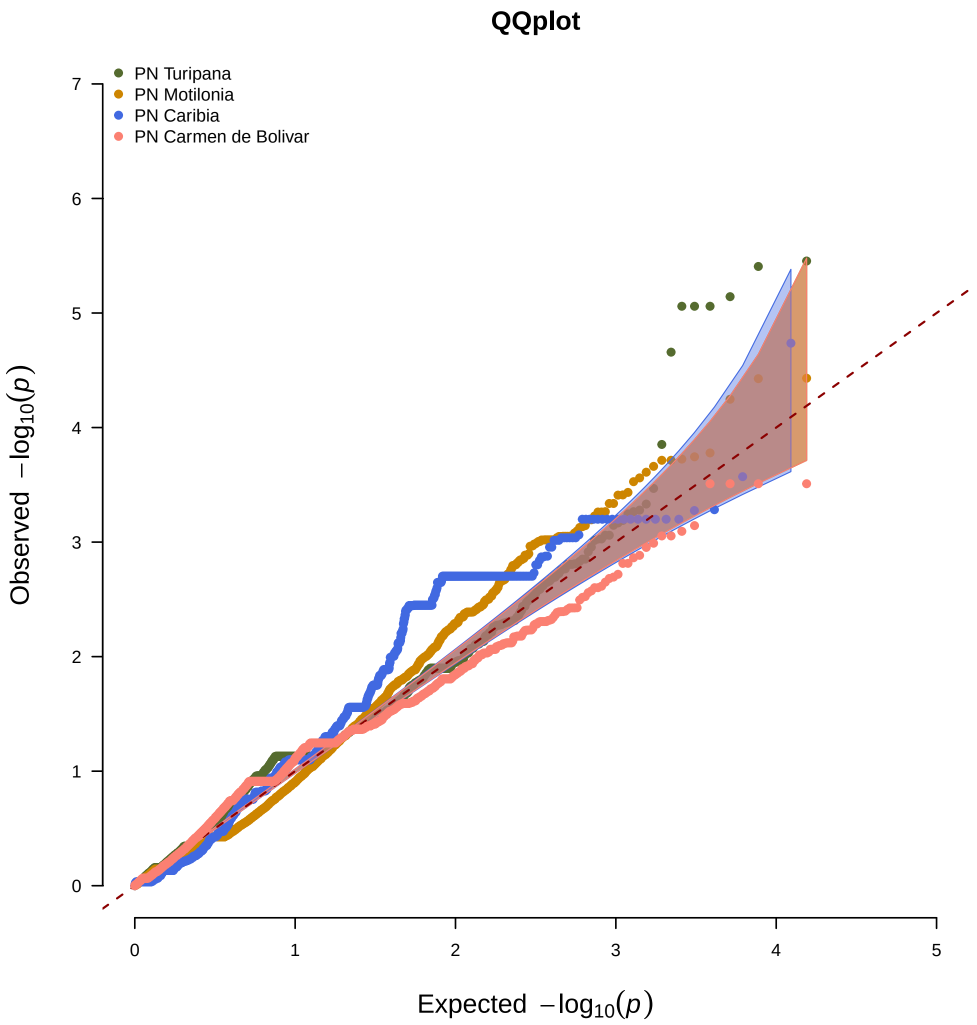

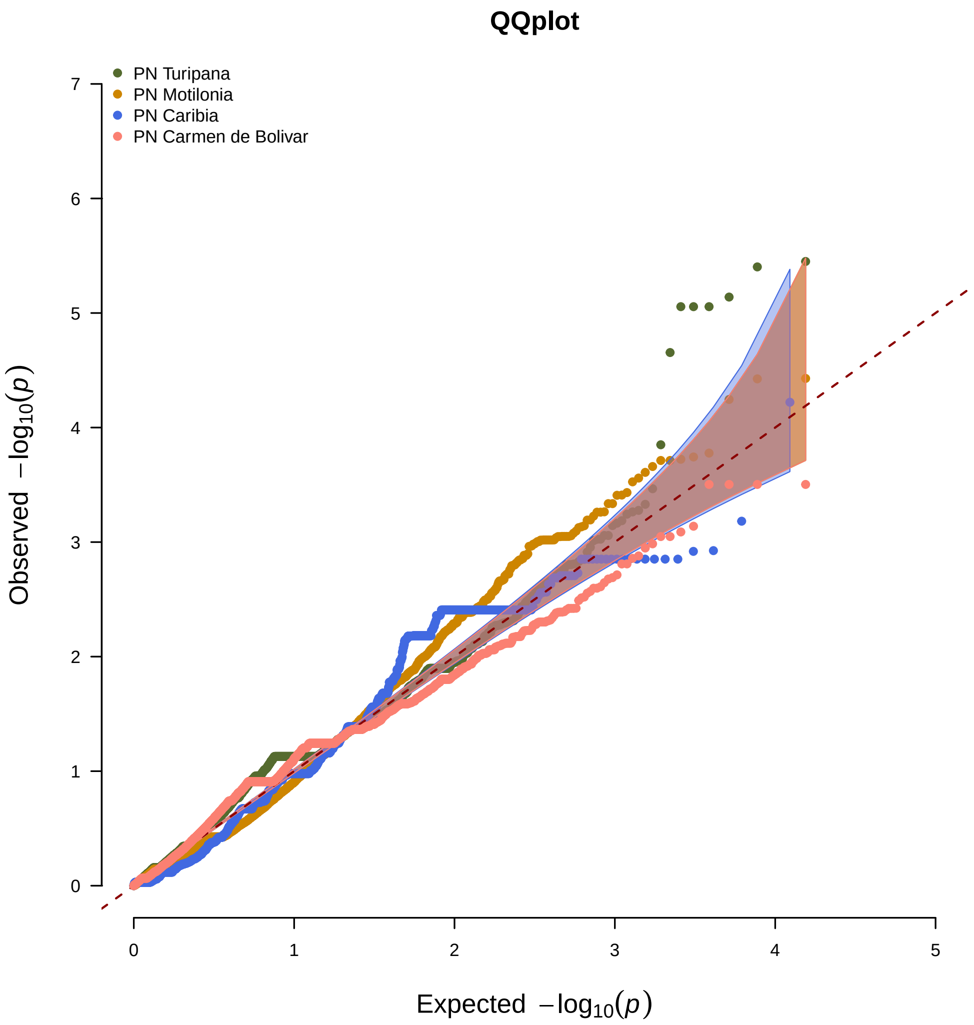

### Supplemental Figure 10

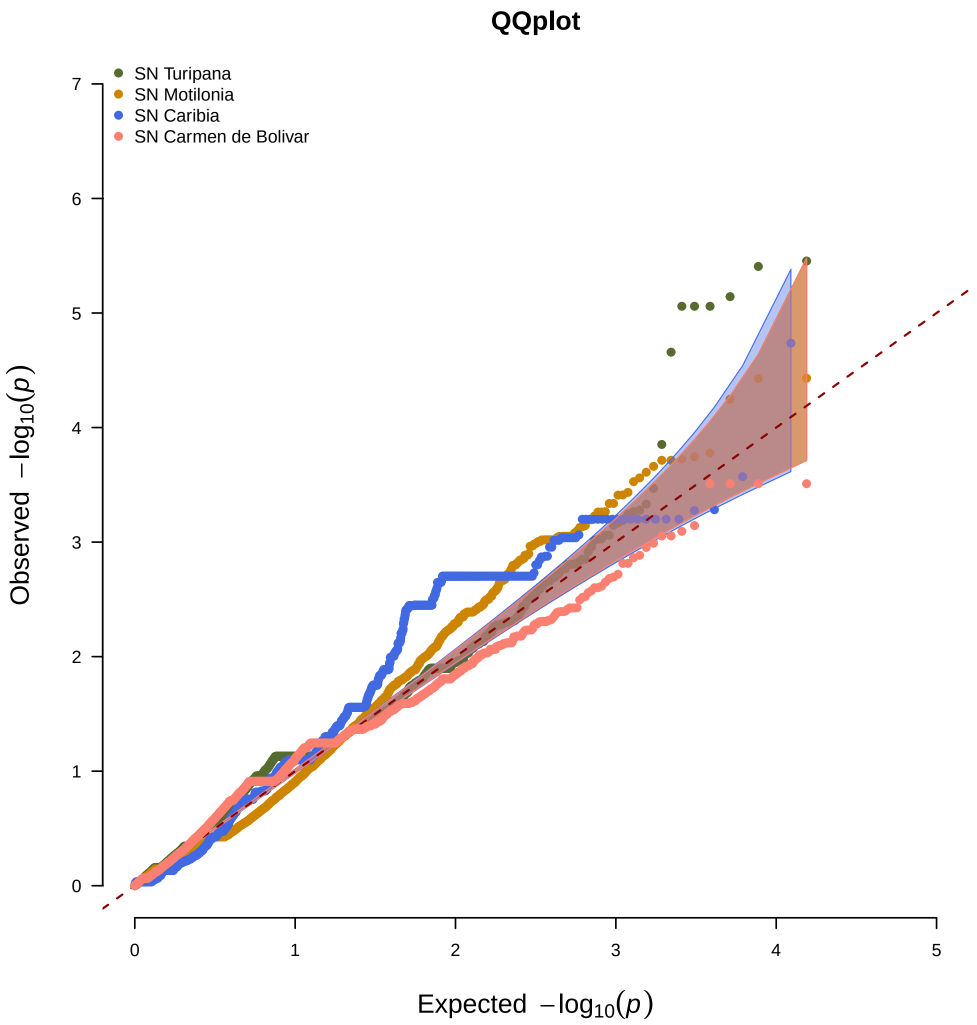

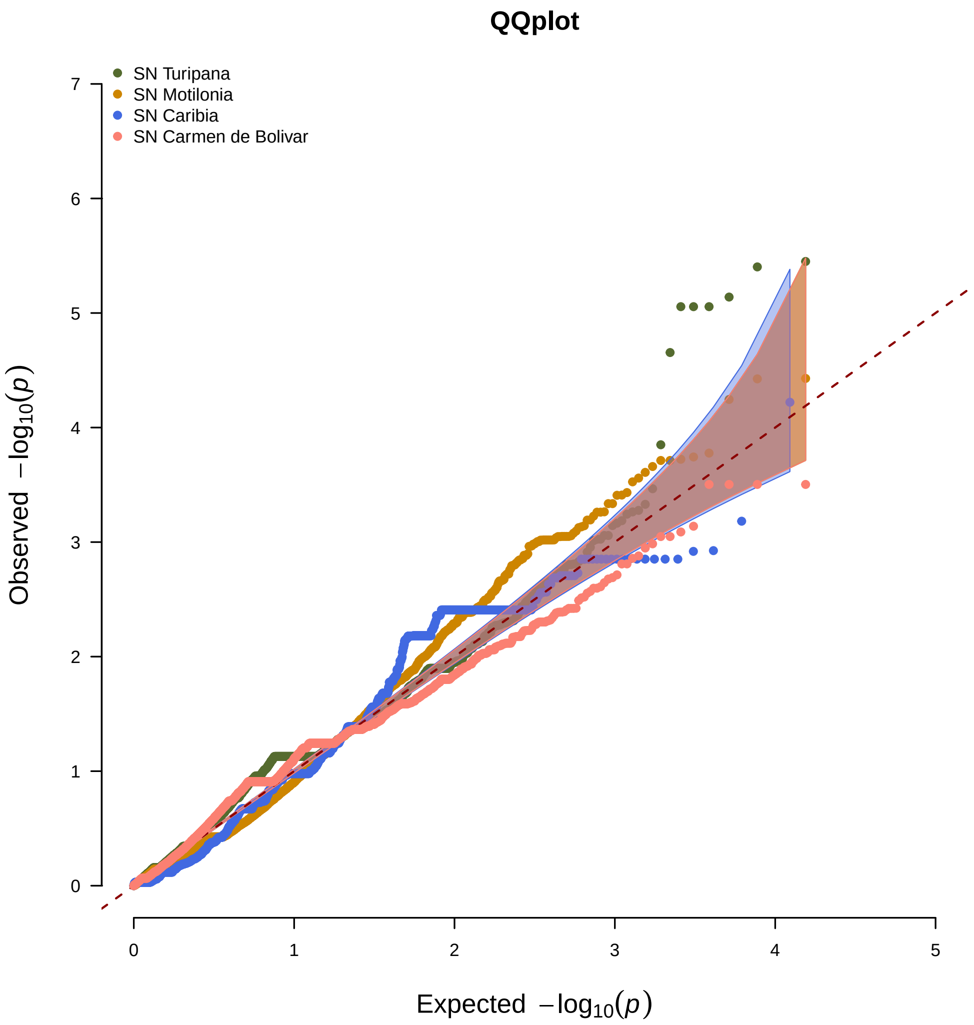

### Supplemental Figure 11

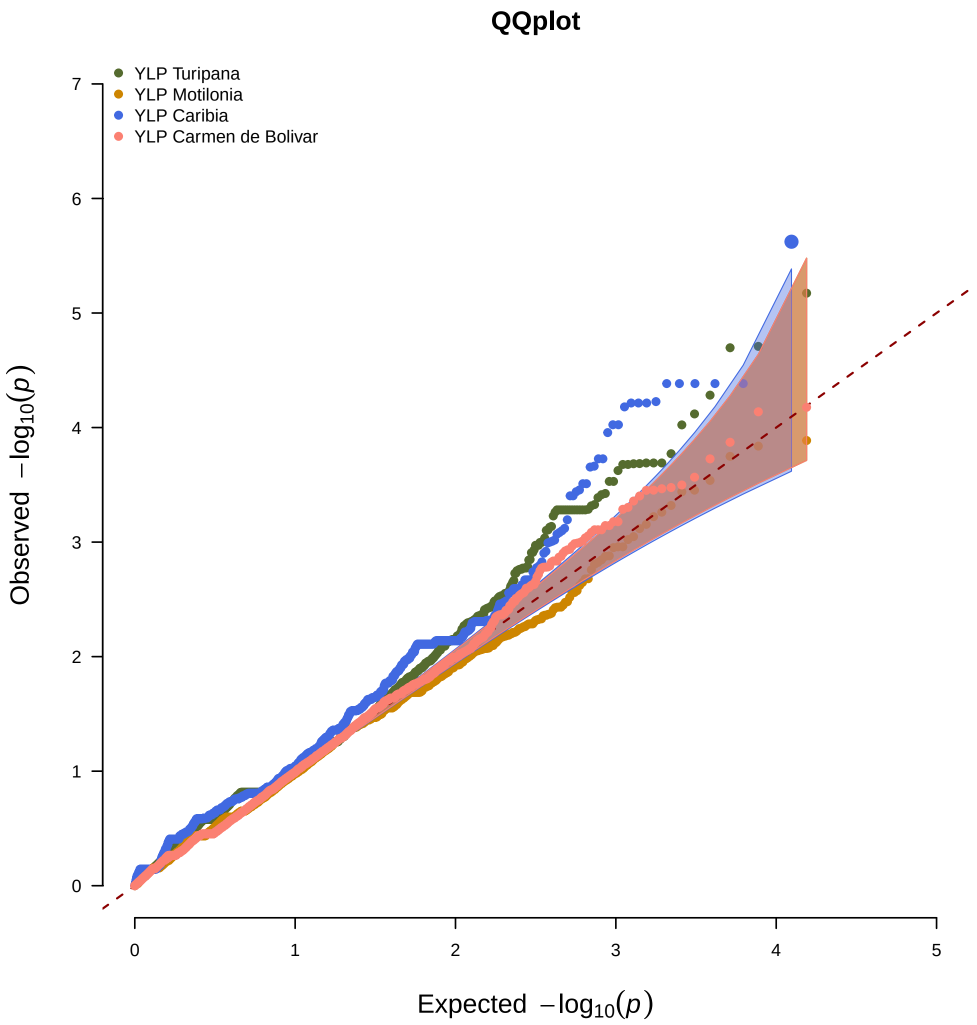

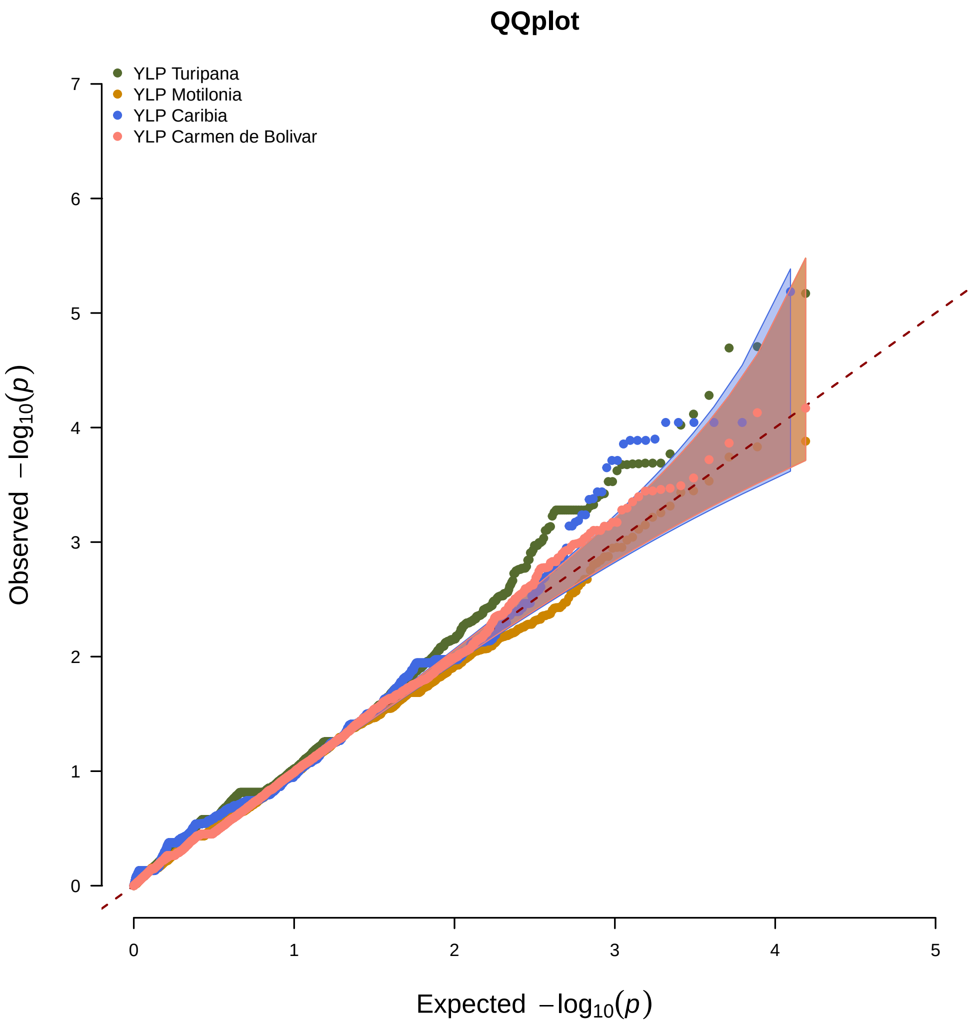

### Supplemental Figure 12

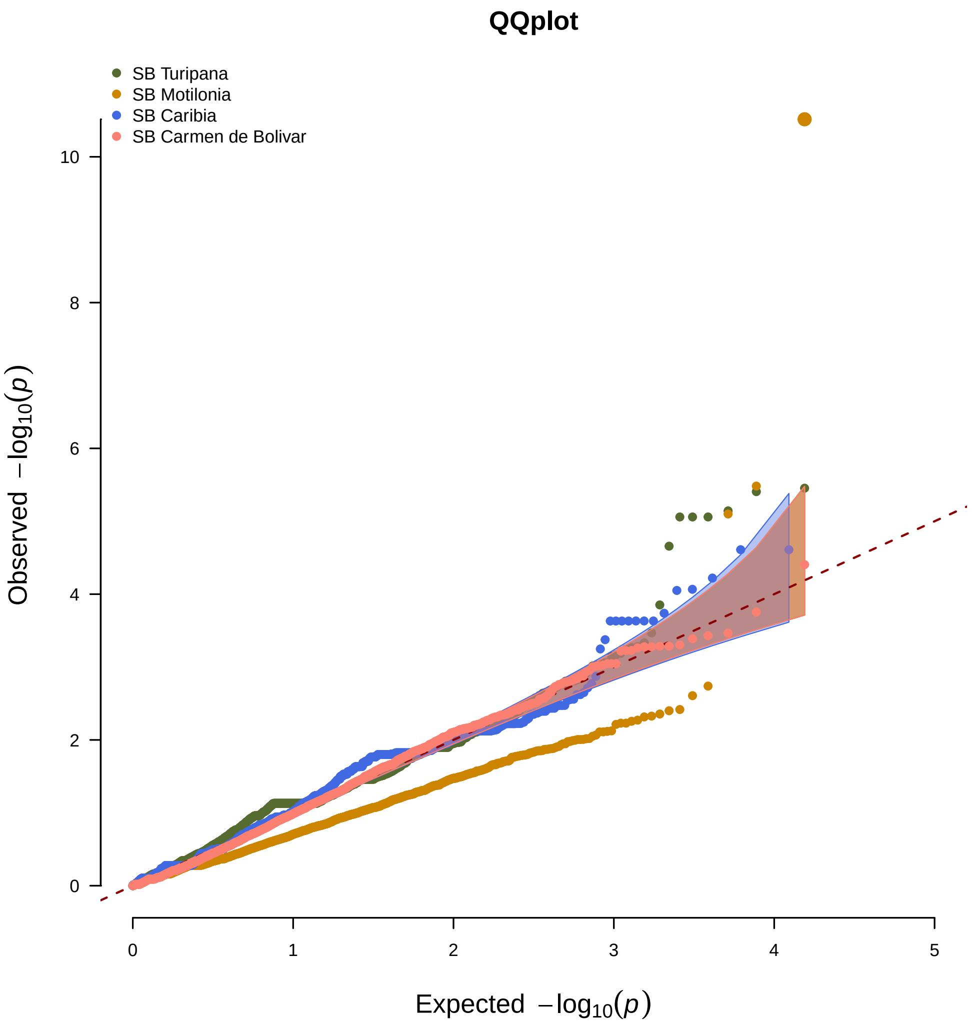

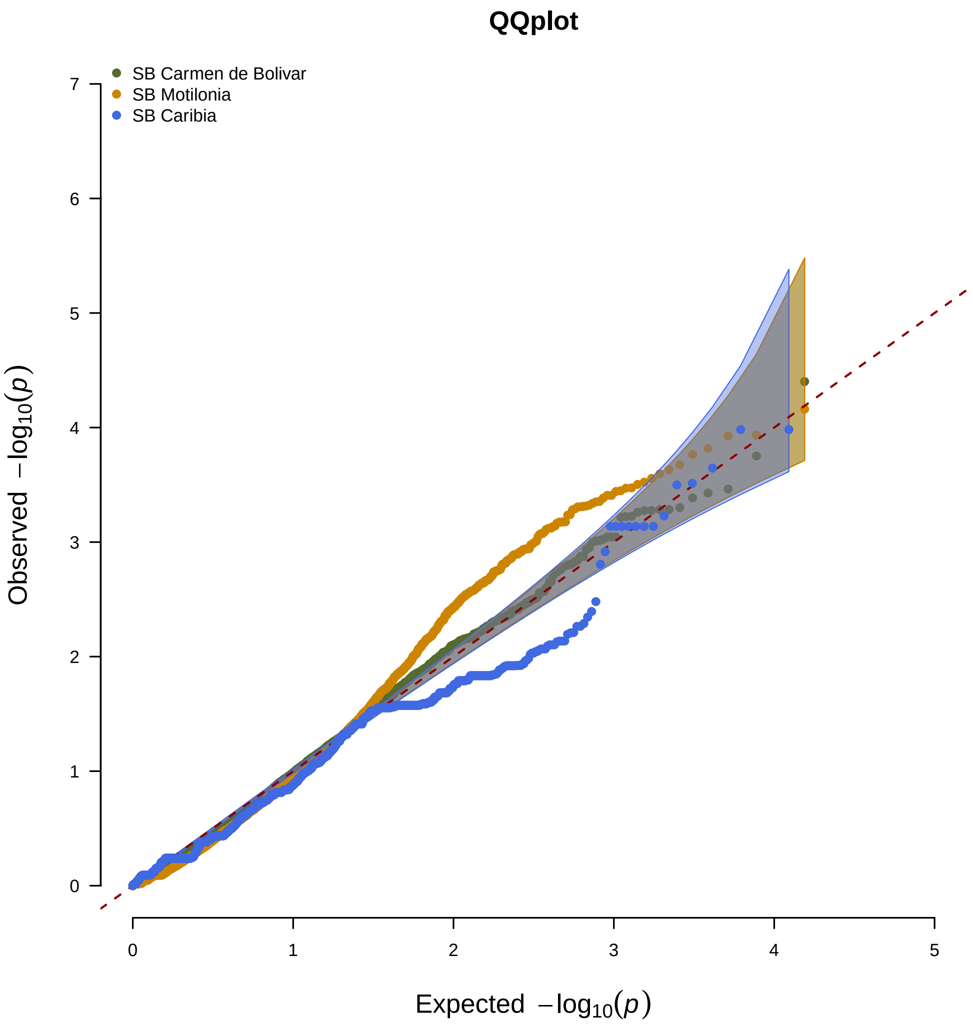

### Supplemental Figure 13

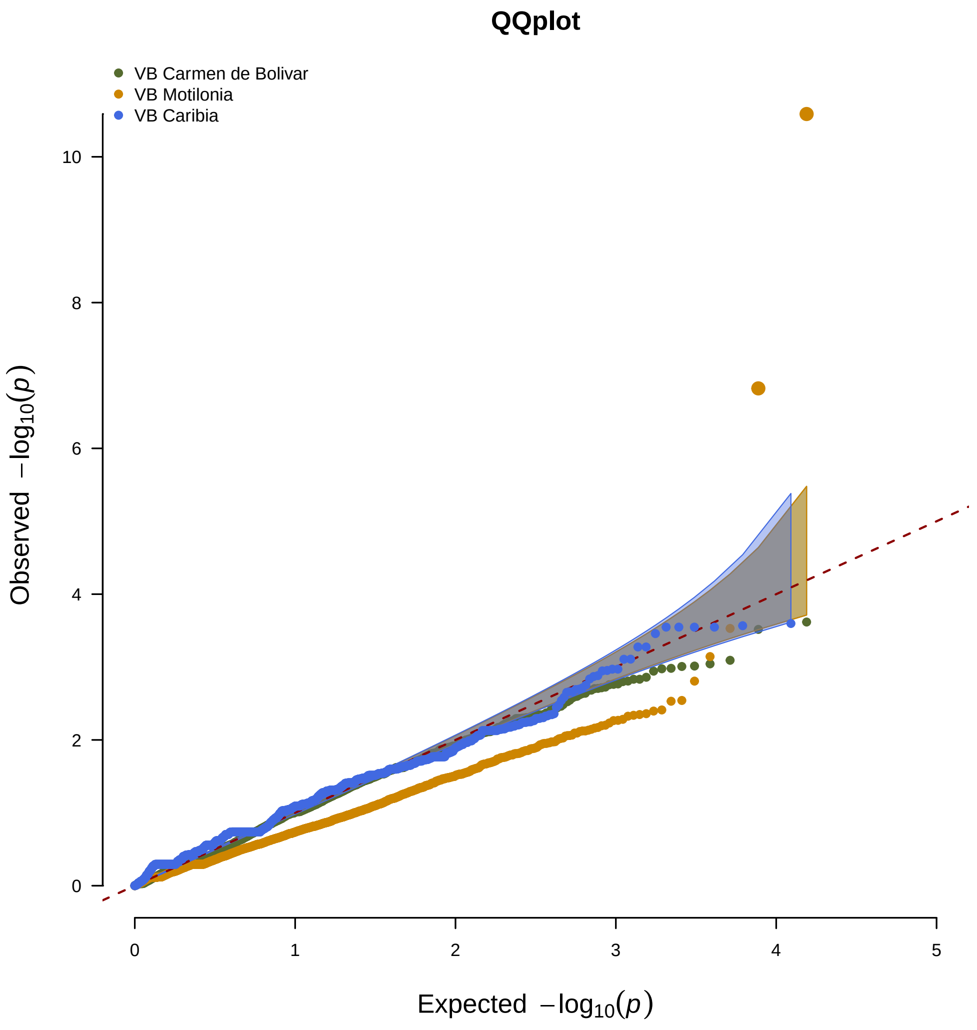

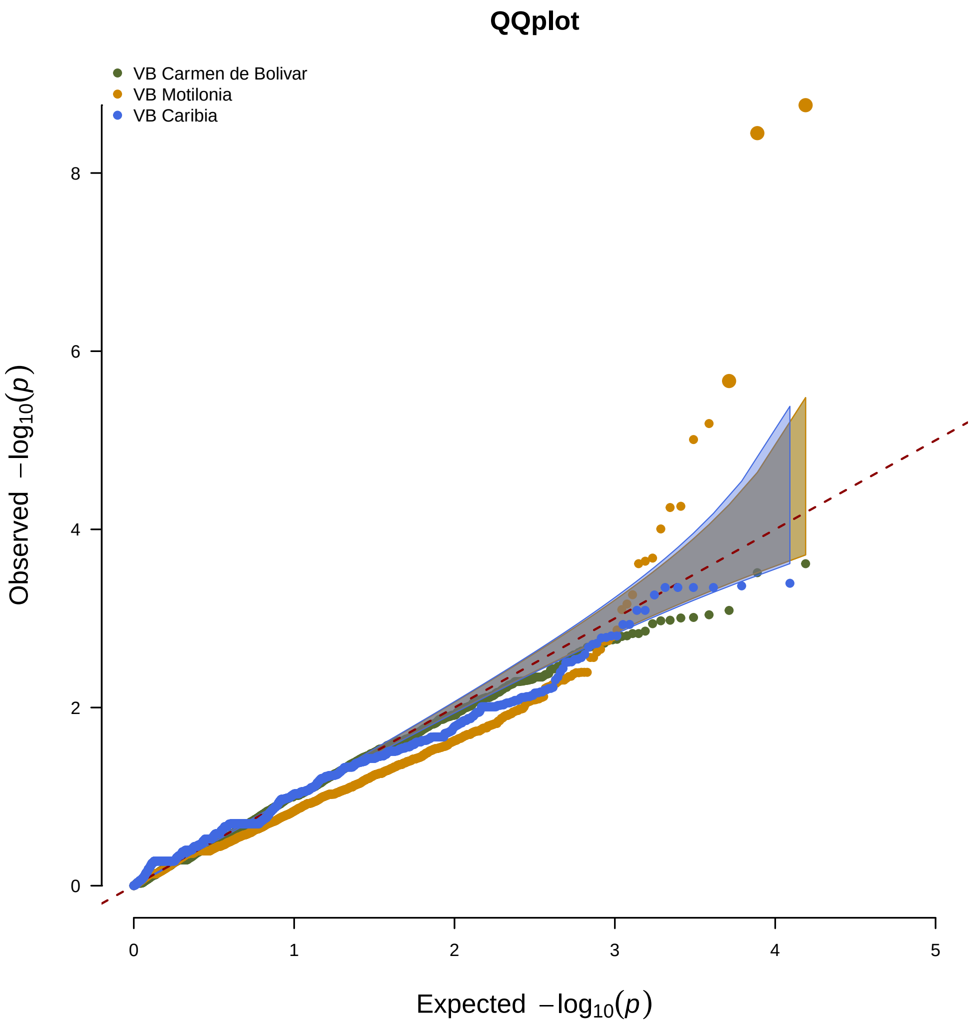
